# A Multicopper Oxidase from *Paenibacillus polyethylenelyticus* JNU01 Oxidizes Polyethylene

**DOI:** 10.64898/2026.02.24.707384

**Authors:** Seung-Do Yun, Seongmin Kim, Seong Jin An, Hyun-Woo Kim, Jin-Hee Cho, Hyeoncheol Francis Son, Chul-Ho Yun, Bong Hyun Sung, Gregg T. Beckham, Won Seok Chi, Chungoo Park, Soo-Jin Yeom

## Abstract

Polyethylene (PE) is a widely used plastic that persists in the environment and resists breakdown via microbial degradation. In this work, we discovered a new bacterium, *Paenibacillus polyethylenelyticus* JNU01, that grows on a PE-like wax (PELW, 4 kDa) as its sole carbon source, causing chemical modifications to the substrate and releasing small-molecule products. Genomic and transcriptomic analyses identified a multicopper oxidase (*Pp*MmcO) as a key enzyme candidate for this observed activity. *Pp*MmcO caused surface oxidation, increased hydrophilicity, and the release of oxygenated products such as ketones, alkanes, and alkenoic acids. Scanning electron microscopy confirmed surface damage on both PELW and post-use greenhouse PE films. Weight loss analysis showed mass losses of 5.2% for the PELW powder and 1.6% for the greenhouse PE film after treatment with wild-type *Pp*MmcO. We propose a radical-mediated pathway catalyzed by *Pp*MmcO. These findings identify a new bacterium and enzyme capable of initiating PE oxidation and provide insight into biological processes that may act on polyethylene.

## 1. Introduction

The management of plastic waste is recognized as a clear, global challenge ^1–3^. Polyethylene (PE), which accounts for approximately 30% of global plastic production, has a long lifetime in the biosphere and tends to accumulate in both marine and terrestrial environments, making it a major contributor to plastic pollution ^2, 4–6^. Given the long predicted environmental lifetime of PE and its chemical stability, it is intriguing to consider whether microorganisms have evolved, or harbor sufficiently promiscuous catabolic pathways, to initiate its degradation ^6^.

For many decades, researchers have sought microbes capable of PE degradation ^7^, and numerous studies have hypothesized that the enzymatic mechanisms involved in PE degradation likely include radical-mediated, oxidative reactions. Specifically, it is hypothesized that introducing oxygen into the stable and inert C−C backbone of PE is crucial, resulting initially in the formation of carbonyl groups ^6, 8^. Subsequently, these oxidized sites could facilitate the cleavage of long hydrocarbon chains to smaller molecules ^7, 9^. Several oxidative enzymes have been implicated in PE degradation, including alkane monooxygenase (AlkB) ^10^, cytochrome P450 ^11, 12^, peroxidase ^13^, laccase ^13, 14^, and laccase-like multicopper oxidases ^15, 16^, have been reported to oxidize the PE and promote its degradation. Despite these promising findings, the enzymatic mechanisms underlying PE degradation remain unclear as to their relevance or even existence for realistic, environmental PE breakdown ^7^. However, recent advances in environmental microbiology, genomics, and polymer characterization offer opportunities to identify enzymes capable of oxidative PE degradation. At the same time, it is important to note that the term “biodegradation” is often used inconsistently in the literature, and complete mineralization of PE requires isotope-labeled polymers and carbon-balance measurements, which remain largely absent in the field^17^. Accordingly, this study focuses on oxidative transformations of PE rather than implying full biodegradation.

In this work, we isolated a bacterium, *Paenibacillus polyethylenelyticus* JNU01, which grew in minimal medium containing a PE-like wax (PELW) substrate – the commonly used 4 kDa “PE” from Sigma-Aldrich – as the sole carbon source and demonstrated its partial conversion. To identify putative enzymes responsible for this microbial growth, whole-genome sequencing and transcriptomics were conducted, revealing an up-regulated multicopper oxidase (MmcO) that was functionally expressed in *Escherichia coli* and biochemically characterized. The purified *Pp*MmcO^WT^ and its variant *Pp*MmcO^H499A^ changed the chemical structure of PELW powder, indicating oxidative modification of the substrate. Small molecule products were detected from *in vitro* assays with the PELW substrate, including ketones, alkanes, and alkenoic acids, showing the ability of *Pp*MmcO to oxidize this substrate. Scanning electron microscopy revealed surface damage on both the solution-cast PELW film and post-use greenhouse PE film following enzymatic reactions with *Pp*MmcO^WT^ and *Pp*MmcO^H499A^. Weight loss analysis showed mass losses of 5.2 ± 0.3% and 5.0 ± 1.8% for the solution-cast PELW film after treatment with *Pp*MmcO^WT^ and *Pp*MmcO^H499A^, respectively, and losses of 1.6± 0% for the post-use greenhouse PE film. Overall, this study identifies *Pp*MmcO as a biocatalyst for partial oxidation of a PE model substrate, providing insights into enzymatic mechanisms for attacking C–C-linked polymers.

## 2.1. Results

### 2.1. Isolation and growth curve of PELW-active bacteria

A low molar mass PELW substrate in powder form (Mw ≈ 4,000, Mn ≈ 1,700 by GPC) was reprecipitated to obtain a powder with an average particle size 100–300 µm, aimed at increasing the external surface area, and GPC analysis confirmed that its molecular weights remained unchanged after reprecipitation. A microbial strain capable of growing on PELW as the sole carbon source was isolated from landfill soil samples collected in Gwangju, Republic of Korea (Figure S1A). On day 40 of the screening, the OD_600_ fold-change results for strain 2GT37 showed an approximate 2.67-fold increase compared to day 0 (Figure S1B). A phylogenetic tree was constructed using 22 16S rRNA sequences, identifying the strain as belonging to the genus *Paenibacillus* (Figure S1C). Thus, 2GT37 was initially designated as *Paenibacillus* sp. JNU01.

To characterize *Paenibacillus* sp. JNU01’s growth at various PELW concentrations, the growth curve of the strain was measured in M9-TES medium containing three PELW concentrations: 10, 30, and 50 mg/mL. Flask cultures were incubated in shaking incubators (28 °C, 200 rpm), ensuring sufficient aeration (see Methods). Growth increased progressively during the first 20 days of the culture, after which it plateaued, indicating entry into stationary phase. On day 20, the OD_600_ value reached 0.44 at 10 mg/mL PELW, 1.33 at 30 mg/mL PELW, and 1.85 at 50 mg/mL PELW (Figure 1A). In addition, the turbidity of the media varied with PELW concentration, and we confirmed that the microorganisms rendered the PELW powder hydrophilic, causing it to become suspended in and distributed throughout the media (Figure 1A), suggesting a modification in the surface properties of PELW ^8, 11^.

**Figure 1.**
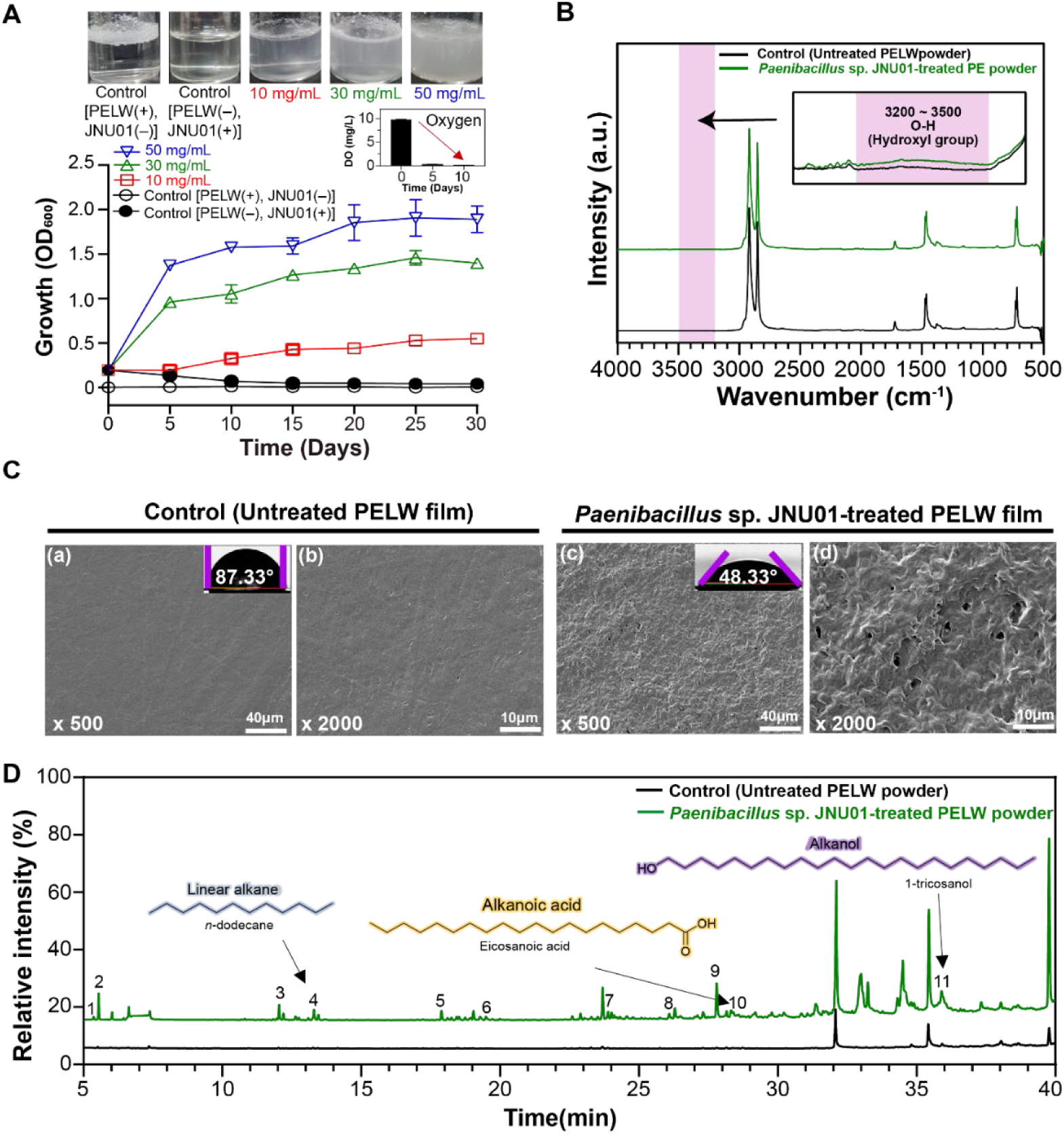
Analysis of the PELW activity of *Paenibacillus* sp. JNU01. (**A**) The growth curves (*n* = 2, independent experiments) of *Paenibacillus* sp. JNU01 at various PELW concentrations (10, 30, and 50 mg/mL). The inset image shows the dissolved oxygen (DO) analysis results. (**B**) FT-IR spectra of control (untreated PELW powder) and *Paenibacillus* sp. JNU01*-*treated PELW powder. (**C**) Top-view SEM images of control (a and b; untreated PELW film) and *Paenibacillus* sp. JNU01-treated PELW film (c and d). Inset images show the corresponding average water contact angle analysis results. (**D**) GC-MS analysis results from the reaction media of the control (untreated PELW powder) and *Paenibacillus* sp. JNU01-treated PELW powder.

To further elucidate the strain’s activity during growth on PELW, we conducted a dissolved oxygen (DO) analysis. While the initial DO level was 9.82 mg/L, the DO level decreased to 0.255 mg/L on day 5 and to 0.155 mg/L on day 10 (inset image in Figure 1A). Together, these results indicate that oxygen consumption occurs during the microbial growth on PELW ^18^.

### 2.2. Chemical and physical changes to PELW during growth of *Paenibacillus* sp. JNU01

Fourier transform infrared (FT-IR) spectroscopy was initially used to analyze the chemical modifications of the post-digestion PELW samples treated with *Paenibacillus* sp. JNU01. However, as recently emphasized by Stepnov *et al.* ^19^, biomolecule contamination, particularly from residual proteins or microbial debris, can lead to misleading FT-IR signals in PE samples after biological treatment. To address this concern, we tested extensive washing protocols to determine whether rigorous cleaning could eliminate false-positive signals originating from biomolecular residues. Comparison of FT-IR spectra before and after extensive washing revealed that thorough washing is essential for reliable data interpretation. Specifically, peaks at 1,252 cm^-1^ (peak 1), 1,540 cm^-1^ (peak 2), and 1,650 cm^-1^ (peak 3), corresponding to amide-related vibrations, disappeared after sequential washing with methanol (4 times), 1 M NaOH (5 times), and distilled water (5 times), confirming that these signals arose from residual biomolecules (Figure S2). These results highlight the necessity of thorough sample pretreatment prior to FT-IR analysis to ensure that observed spectra changes truly reflect chemical modifications in PELW. This careful approach is particularly important given recent discussions emphasizing that oxidation-related spectral features in polyethylene should not be overinterpreted^17^.

After applying this extensive washing protocol, the control (untreated PELW powder) showed FT-IR spectra consisting mainly of hydrocarbons with C–C and C–H bonds, exhibiting strong absorption bands at 1,480 cm^-1^ and 2,900 cm^-1^ ^20, 21^. In contrast, the *Paenibacillus* sp. JNU01-treated samples exhibited small spectral changes at 3,200–3,500 cm^-1^ (O–H stretching) ^20, 22, 23^ indicating the formation of oxygen-containing functional groups such as hydroxyl (Figure 1B). These spectral results suggest that *Paenibacillus* sp. JNU01 treatment may facilitate the incorporation of oxygen-containing functional groups into the hydrocarbon backbone of PELW. However, because the FT-IR peak changes were relatively minor, additional analyses were conducted to further verify the activity of *Paenibacillus* sp. JNU01.

The physical change to PELW film surfaces during cultivation of *Paenibacillus* sp. JNU01 was assessed using scanning electron microscopy (SEM) and average water contact angle values. Control (untreated PE film) showed a smooth and dense surface morphology without significant defects (Figure 1C(a), (b)), while the *Paenibacillus* sp. JNU01-treated PELW film exhibited a noticeably porous structure, with pores up to a few microns in size, and a rough surface texture, suggesting modification of PELW polymer chains during bacterial growth (Figure 1C(c), (d)). In addition, the average water contact angle of the control (untreated PELW film) was 87.33° (inset image in Figure 1C(a)). In contrast, the *Paenibacillus* sp. JNU01-treated PELW film exhibited a reduced water contact angle of 48.33° (inset image in Figure 1C(c)). This result indicates that *Paenibacillus* sp. JNU01-induced PELW film activity leads to a relatively hydrophilic surface, possibly attributable to oxidation.

### 2.3. Metabolites in the PELW media

To identify the PELW metabolites released in the PELW media inoculated with *Paenibacillus* sp. JNU01, a gas chromatography-mass spectrometry (GC-MS) analysis was performed, confirming the production of alkanes, alcohols, and carboxylic acids (Figure 1D and Table S1). The organic compounds matched with the NIST/WILEY library as 3,4,5-trimethylheptane, 2,3,3-trimethylheptane, 3-ethyl-3-methylheptane, *n*-dodecane, 5-methyl-5-propylnonane, *n*-tetradecane, *n*-hexadecane, 11-oxododecanoic acid, *n*-eicosane, eicosanoic acid, and 1-tricosanol. Linear alkanes, alkanols, and alkanoic acids are potential intermediates from bacterial enzymatic activity during growth on PELW, further suggesting that *Paenibacillus* sp. JNU01 can modify PELW polymers and release small-molecule products.

### 2.4. Discovery of upregulated PELW-active enzyme in *Paenibacillus* sp. JNU01

First, we compared the *Paenibacillus* sp. JNU01 genome sequence to other bacterial genomes to determine the average nucleotide identity (ANI) for taxonomic purposes. *Paenibacillus* sp. JNU01 exhibited 91.4% ANI with *P. amylolyticus*, covering 83.1% of the *Paenibacillus* sp. JNU01 genome (Supplementary data S1). However, due to the low coverage, which posed challenges for accurate identification, we renamed strain JNU01 to *Paenibacillus polyethylenelyticus* JNU01. To elucidate potential enzymes that might act on PELW, we conducted RNA-Seq to examine the transcriptional response of *P. polyethylenelyticus* JNU01 to PELW exposure. Since the genome was first sequenced and analyzed (Figure 2A), a reference-based RNA-Seq approach, which offers higher accuracy in gene expression analysis compared to *de novo* approaches ^24^, was applied. The RNA-Seq analysis was performed under two independent experimental conditions: culturing in M9-TES medium supplemented with glucose (denoted as the “Glucose” condition), with four biological replicates, and culturing in M9-TES medium supplemented with PELW (denoted as the “PELW” condition), with two biological replicates. From these experiments, an average of 24.4 million raw sequencing reads were generated (Supplementary data S2). After preprocessing, an average of 15.9 million high-quality, clean reads were uniquely mapped to the newly sequenced *P. polyethylenelyticus* JNU01 genome, achieving an average mapping rate of 94.3%. In total, 1,141 differentially-expressed enzymes (DEEs) were identified, including 343 upregulated and 798 downregulated enzymes in the PELW condition compared to the glucose condition (Figure 2B and Supplementary data S3 and S4).

**Figure 2.**
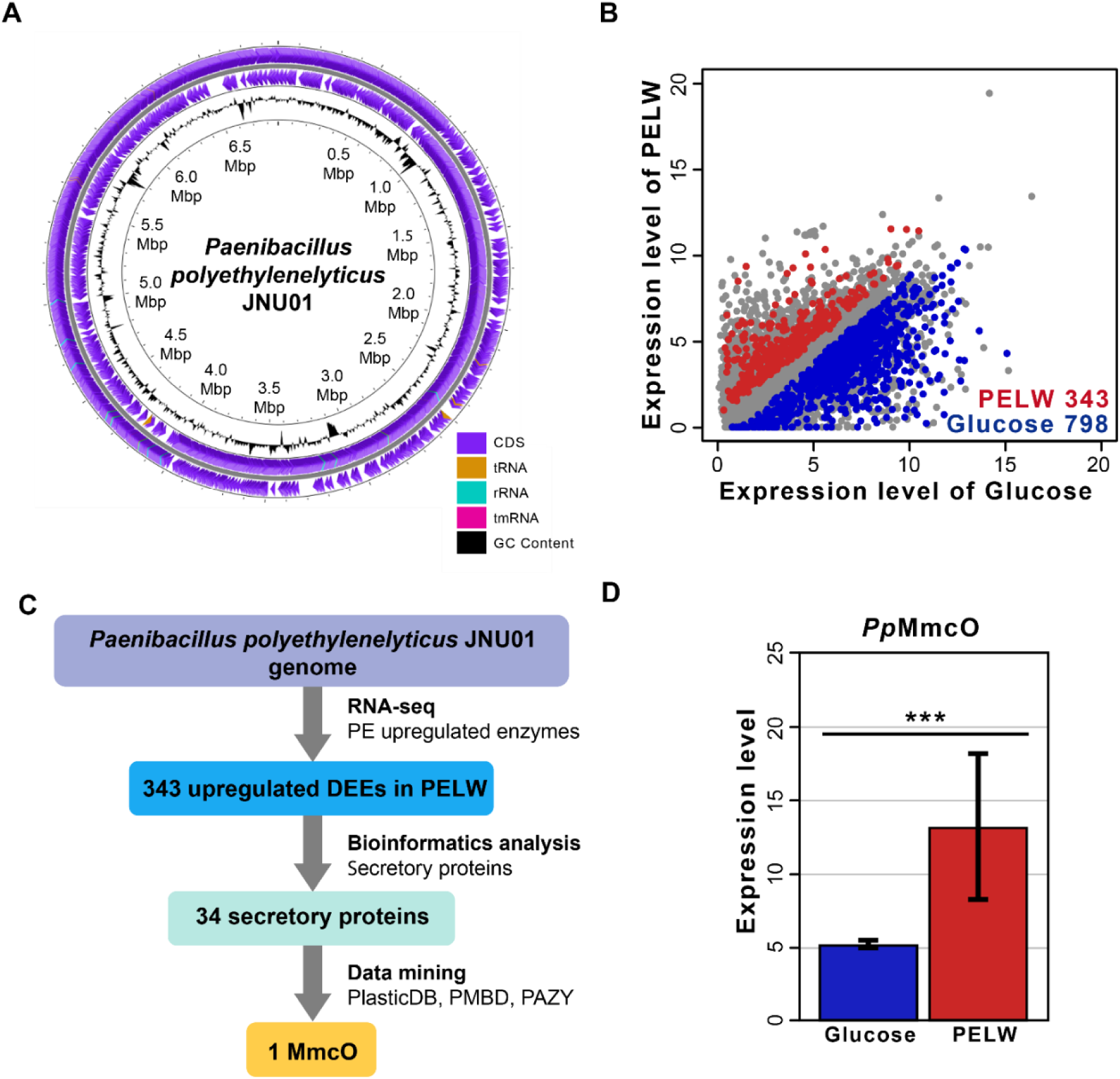
Genome and transcriptome analysis of *P*. *polyethylenelyticus* JNU01. (**A**) A Circos diagram of the genome: (from the outer to inner rings) 1) forward strand features, 2) reverse strand features, and 3) GC content. (**B**) A scatter plot comparing the transcriptome profiles after culturing with glucose or PELW. Expression levels are shown as log_2_-transformed counts per-million mapped reads. Red and blue points indicate significantly-upregulated genes in PELW and glucose cultures, respectively. (**C**) The strategy for identifying potential PELW-degrading enzymes among the 343 genes upregulated in PELW cultures. (**D**) The expression levels of NLBNIHML_00764 (multicopper oxidase *Pp*MmcO); *p*-values were calculated using edgeR, and adjusted *p*-values (FDR) are shown. Error bars represent the standard error of mean, and asterisks (***) indicate significant differences at *p* < 5 × 10^−7^.

Cascade enzymatic reactions such as those involving alkane hydroxylation (e.g., P450s, UPOs, and AlkB), alcohol dehydrogenase (Adh), aldehyde dehydrogenase (Aldh), Baeyer–Villiger monooxygenase (BVMO), and esterase have been proposed as potential pathways for PE degradation ^8^. Also, a recent study further demonstrated that pre-oxidized PE fragments can undergo chemo-enzymatic conversion via catalase–peroxidase, ADH, BVMO, and lipase ^25^. Transcriptomic analysis of *P. polyethylenelyticus* JNU01 revealed no upregulation of alkane hydroxylation–related enzymes (e.g., P450s, UPOs, and AlkB) under the PELW conditions, suggesting that this strain may employ an alternative oxidative mechanism for PELW degradation. Given these findings, it was hypothesized that alternative enzyme candidates for PELW degradation likely exist and are potentially secreted extracellularly and upregulated in response to PELW exposure. Using SignalP 6.0, 34 of the 343 upregulated proteins were predicted to be secreted enzymes (Figure 2C), and a diverse set of hydrolases, oxidoreductases, lyases, and a nuclease was identified, supporting the presence of multiple enzymatic functions potentially involved in PELW activity. Detailed annotations of these enzymes are provided in Supplementary data S3.

By cross-referencing with enzyme databases (PlasticDB, PMBD, and PAZY)^26–28^, we identified a single enzyme, upregulated in presence of PELW, MmcO (multicopper oxidase), that was hypothesized to be a key enzyme for PE activity (Figure 2D). The enzyme was named *Pp*MmcO following its identification as a multicopper oxidase from *P. polyethylenelyticus* JNU01, based on its genetic and functional characteristics. Previous research has shown that mixed-function oxidases tend to hydroxylate linear alkanes and fatty acids ^29^, while laccase-like multicopper oxidases have been suggested as potential enzymes for PE oxidation ^15, 16^. Amino acid sequence alignments performed using SnapGene (GLS Biotech, San Diego, CA, USA) showed that *Pp*MmcO exhibited low identities of 30.59% and 23.37% with MCOs from *Rhodococcus opacus* R7 ^15^ and *Klebsiella pneumoniae* Mk-1 ^16^, respectively. Additionally, upregulated genes involved in Adh (one enzyme), Aldh (two enzymes), esterase (seventeen enzymes), and lipase (two enzymes) (Figure S3), which might participate in further degradation processes of hydroxylated-PE ^8^, were identified. Signal peptide prediction (SignalP) revealed that the gene sequences encoding two esterases contained signal peptide regions, whereas those encoding the other enzymes were not predicted to contain such sequences. These findings suggest that *Pp*MmcO, along with the other identified enzymes, may be part of a coordinated system for PE oxidation, with *Pp*MmcO potentially serving as a key secreted oxidase enzyme.

To further study the function of *Pp*MmcO through evolutionary relationships, a phylogenetic analysis was conducted. The MCO family encompasses diverse proteins, including laccases, ascorbate oxidases, and ferroxidases ^30^. The *Pp*MmcO amino acid sequence was aligned with those of MCOs with known enzyme commission (EC) numbers and reported PE oxidation activity. This alignment served as the foundation for constructing a phylogenetic tree (Figure S4 and Supplementary data S5) that included a wide range of MCO types. The analysis revealed clear clustering patterns among different MCO families. Notably, *Pp*MmcO was found to cluster within the laccase clade, strongly suggesting its functional identity as a laccase enzyme.

### 2.5. Optimization of catalytic activity in the wild-type *Pp*MmcO

*Pp*MmcO had very low sequence identities compared to other MCOs ^15, 16^ and contains a putative signal peptide identified by SignalP (v6.0) ^31^ (Table S3). A signal peptide-deleted recombinant wild-type *Pp*MmcO (*Pp*MmcO^WT^) was purified, with the SDS-PAGE results showing a molar mass of 60 kDa (Figure S5). Subsequently, the catalytic activity of *Pp*MmcO was investigated using an oxidation assay using 2,2’-azino-bis(3-ethylbenzothiazoline-6-sulfonic acid (ABTS) as a substrate. The optimal conditions for *Pp*MmcO^WT^ activity were at 65 °C and a pH of 3.6 in a 2.0 mM copper ion medium (Figure S6A-C). The relative activity of *Pp*MmcO^WT^ increased with temperature, peaking at 65 °C, and then sharply decreased at 70 °C and above. The *Pp*MmcO^WT^ exhibited the highest activity in Na-acetate buffer (pH 3.6), and its activity plateaued and became saturated at a 2.0 mM Cu^2+^. The activity of *Pp*MmcO^WT^ with 2.0 mM Cu^2+^ was 4.45-fold higher than without copper. In 5-day thermal stability tests, 50% residual activity was observed after 6 hours at 65 °C, 1 day at 50 °C, and 4 days at 30 °C (Figure S6D). Additionally, *Pp*MmcO activity was evaluated under near-neutral pH conditions using phenolic substrates such as 2,6-dimethoxyphenol (2,6-DMP) and guaiacol. However, in the presence of 2.0 mM CuSO_4_, these substrates underwent rapid non-enzymatic oxidation, making reliable measurements at neutral pH impossible.

### 2.6. PELW powder, solution-cast PELW film, and greenhouse PE film oxidation by *Pp*MmcO

To evaluate the oxidative activity of *Pp*MmcO^WT^ on PELW powder, solution-cast PELW film, and greenhouse-cover PE film, enzyme reactions were conducted at 50 °C for 1 day in 50 mM sodium acetate buffer (pH 3.6) containing 2.0 mM CuSO_4_. For PELW powder, both *Pp*MmcO^WT^ (0.2 mg/mL) and boiled *Pp*MmcO^WT^ (0.2 mg/mL, boiled at 95 °C for 10 min) were tested to verify whether the observed hydrophilization was truly enzyme-dependent, whereas only unboiled *Pp*MmcO^WT^ was used for the PELW film and greenhouse PE film. Both control conditions, untreated PELW powder and PELW powder mixed with *Pp*MmcO^WT^ (boiled at 95 °C for 10 min to remove enzymatic activity), showed no visible mixing in the reaction solution even after 24 h. In contrast, when unboiled *Pp*MmcO^WT^ was added, the PELW powder gradually became suspended with the enzyme solution within 24 h, indicating enzyme-induced hydrophilization of the PELW powder (Figure 3A).

**Figure 3.**
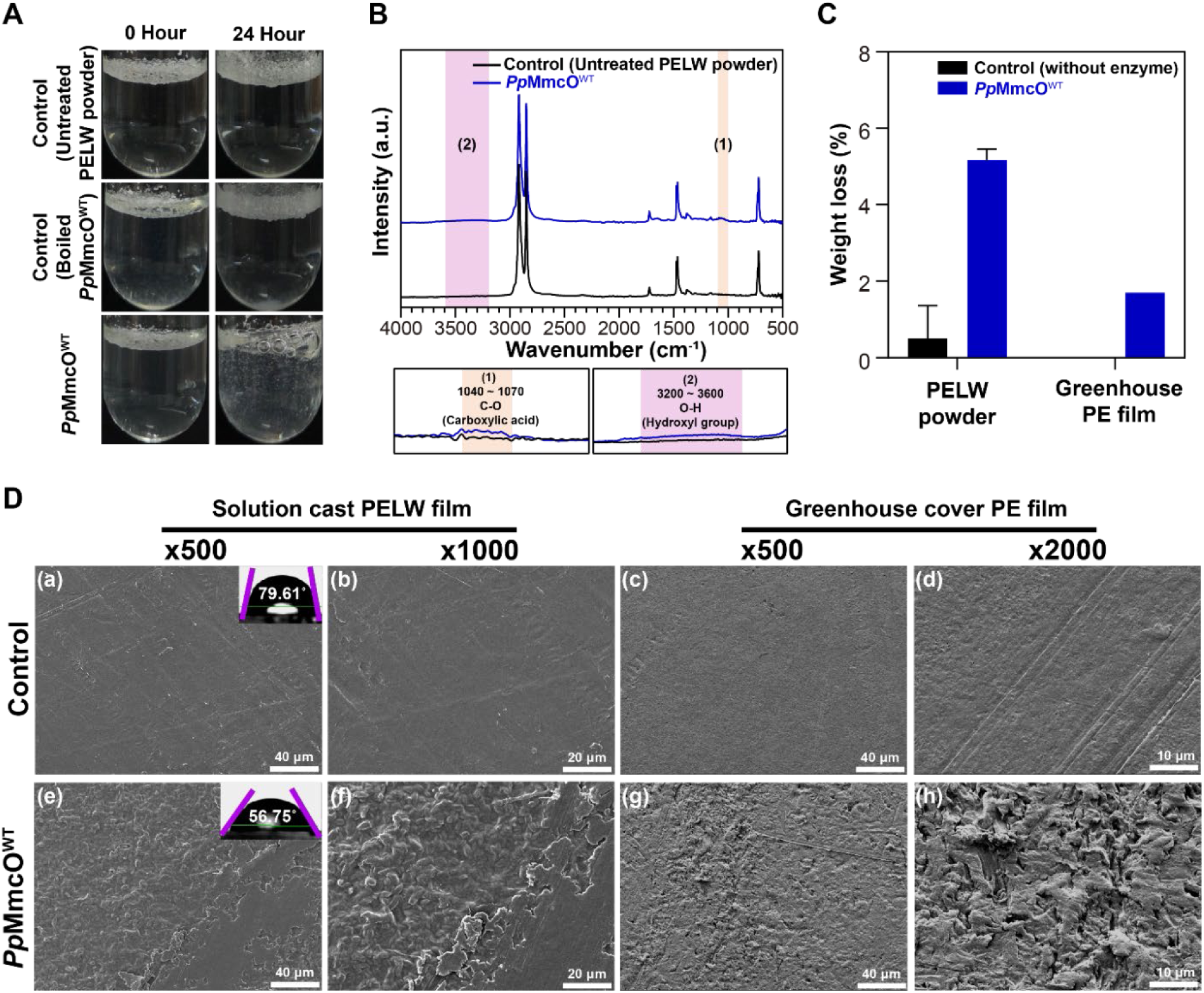
Degradation analysis of PELW powder, solution-cast PELW film, and greenhouse PE film by *Pp*MmcO^WT^. (**A**) Images of the two controls (untreated PELW powder without enzyme or PELW powder with boiled *Pp*MmcO^WT^) and the *Pp*MmcO^WT^ reaction solutions containing PELW powder over time. (**B**) FT-IR spectra of control (untreated PELW powder) and *Pp*MmcO^WT^-treated PELW powder. (**C**) Comparison of the weight loss (%) of PELW powder and greenhouse PE film after treatment with *Pp*MmcO^WT^. Control samples were incubated under identical conditions without enzyme addition (n = 3, mean ± s.d.). (**D**) SEM images of PELW films and greenhouse cover PE films. Solution-cast PELW films (a, b, e, f) and greenhouse cover PE films (c, d, g, h). Panels (a-d) represent untreated control films (without *Pp*MmcO treatment), while panels (e-h) represent *Pp*MmcO^WT^-treated films. The inset images in (a) and (e) show the average water contact angle (WCA) analysis results.

The FT-IR spectra of PELW treated with *Pp*MmcO^WT^ showed new functional group vibration modes, such as C−O stretching (carboxylic acid) and O−H stretching (hydroxyl group), compared to the untreated PE (Figure 3B). These spectral changes support the oxidative modification of PELW by *Pp*MmcO^WT^.

To further confirm enzymatic degradation, weight loss experiments were performed using PELW powder and the post-use greenhouse PE film (Mw = 119,000 and Mn = 18,000) (Figure 3C). Because PELW represents a low molar mass, waxy substrate, additional experiments were conducted to examine whether *Pp*MmcO^WT^ could act on high molar mass PE films. After enzymatic reactions, PELW powder exhibited a weight loss of 5.2 ± 0.3% in the *Pp*MmcO^WT^ treatment, compared with 0.5 ± 0.8% in the control. In the case of the greenhouse PE film, the control sample did not show any weight loss, whereas the *Pp*MmcO^WT^-treated greenhouse PE film showed a mass loss of 1.6 ± 0%. These results demonstrate that *Pp*MmcO^WT^ exhibits oxidative activity not only toward low molar mass PELW but also toward high molar mass PE films.

SEM analyses were also performed on a solution-cast PELW film and a greenhouse-cover PE film to examine whether *Pp*MmcO^WT^ can oxidatively modify industrial PE surfaces. The surface of solution-cast PELW films and greenhouse-cover PE films treated with *Pp*MmcO^WT^ showed noticeable changes relative to those of the untreated PELW and greenhouse-cover PE films (Figure 3D). The solution-cast PELW film and greenhouse-cover PE film no-enzyme control samples exhibited smooth and regular surfaces via SEM (Figure 3D(a-d)). In contrast, *Pp*MmcO^WT^-treated solution-cast PELW film and greenhouse-cover PE films displayed significant surface damage, including roughened textures and visible cracks (Figure 3D(e-h)). The solution cast PELW film treated with *Pp*MmcO^WT^ showed an average contact angle of 56.75° (Figure 3D(e), inset image), which was significantly reduced compared to the untreated control solution-cast PELW film, which showed a contact angle of 79.61° (Figure 3D(a), inset image), indicating the increased hydrophilicity of the treated film surface. Overall, these results highlight the oxidative activity of *Pp*MmcO^WT^, which alters the physical properties of solution cast PELW films and greenhouse PE films and facilitates surface hydrophilization.

Similar to the *in vivo* assays, a GC-MS analysis was performed to identify small-molecule products in the reaction solution containing *Pp*MmcO^WT^ and the PELW control, comparing peaks detected in the experimental group to those detected in the control group (Figure 4A-D, Table S4). Before GC–MS measurement, all samples were derivatized with BSTFA–TMCS. The degradation products matched with the NIST/ Wiley database were 2-heptadecanone, *n*-eicosane, *cis*-10-heptadecenoic acid trimethylsilyl ester, 2-nonadecanone, *trans*-9-octadecenoic acid trimethylsilyl ester, 2-methylpentacosane, *cis*-10-nonadecenoic acid trimethylsilyl ester, 2-nonacosanone, *n*-triacontane, and 2-tritriacontanone, all of which are composed of carbon chains ranging from 17 to 33. The matched products were methyl ketones, alkanes, a 2-methyl alkane, and alkenoic acids. Additionally, Figure 4E shows the relative intensity (%) of each compound type according to the concentration of *Pp*MmcO^WT^, confirming that the production of alkenoic acids increased as the concentration of *Pp*MmcO^WT^ increased. This further highlights the formation of methyl ketones, alkanes, 2-methyl alkane, and alkenoic acids generated during the enzymatic reaction. These results are consistent with the FT-IR results from the *Pp*MmcO^WT^-treated PE powder and the GC-MS analysis profiles of enzyme reaction mixtures containing alkanes, ketones, and carboxylic acids produced by laccase-like multicopper oxidases from *R. opacus* R7 (i.e., LMCO2 and LMCO3) in a previous study ^15^.

**Figure 4.**
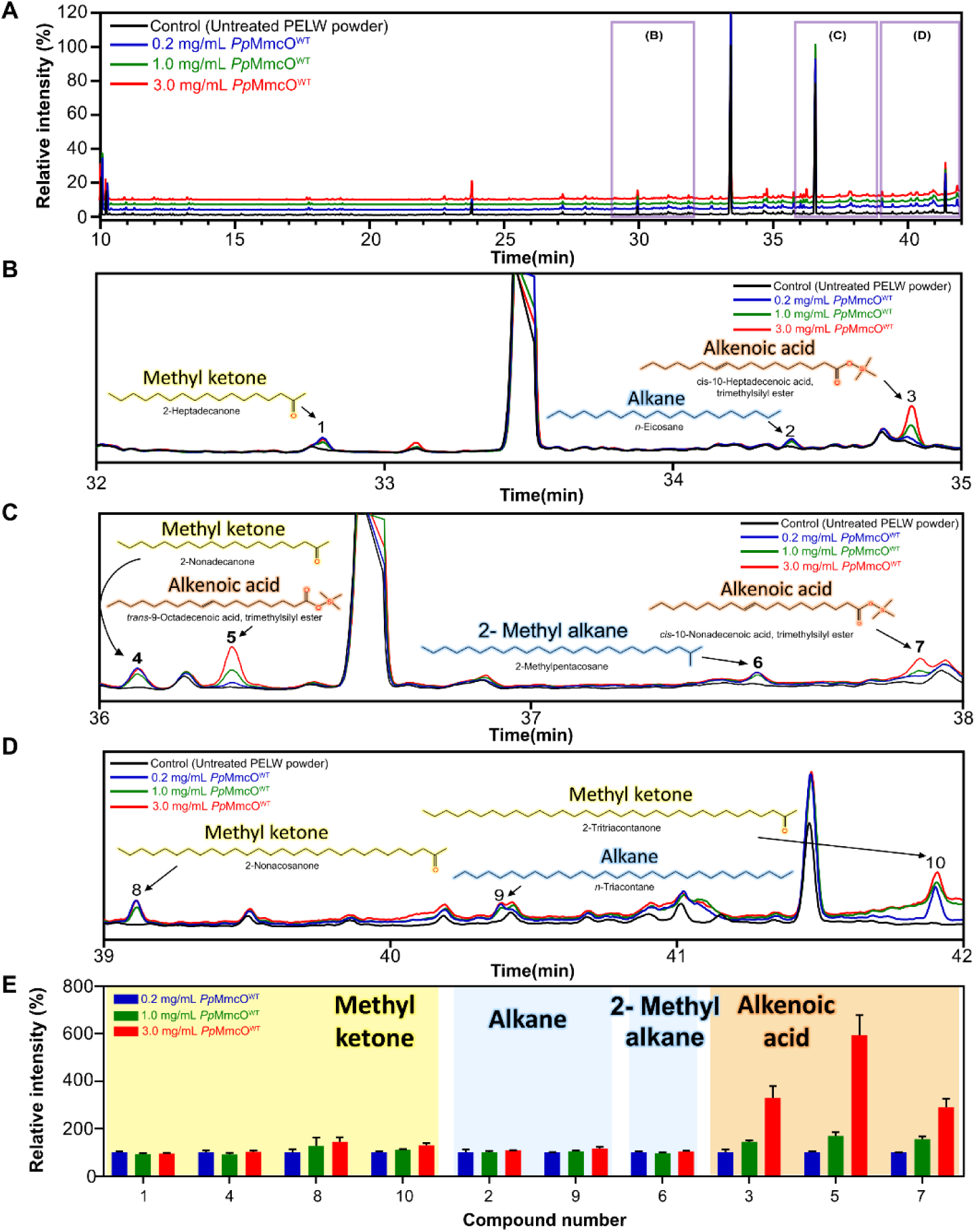
GC-MS analysis of enzyme reaction solutions of *Pp*MmcO^WT^ at different enzyme concentrations. (**A**) GC-MS chromatograms of enzyme reaction solutions with PELW powder treated using different concentrations of *Pp*MmcO^WT^ (0.2, 1.0, and 3.0 mg/mL) compared to those of the control (untreated PELW powder). (**B, C, and D**) Magnified sections of the chromatogram in (A) highlighting detected compounds classified into methyl ketones, alkanes, 2-methyl alkanes, and alkenoic acids. The compounds matched with the NIST/Wiley database are labeled with their respective structures. (**E**) Relative intensity (%) (*n* = 2, independent experiments) of detected compounds grouped by chemical class (methyl ketones, alkanes, 2-methyl alkane, and alkenoic acids) at the different *Pp*MmcO^WT^ concentrations (*n* = 2, independent experiments).

### 2.7. *Pp*MmcO’s structure and the comparative activities of variants

To gain further insights into the *Pp*MmcO enzyme, the structure of *Pp*MmcO was predicted using AlphaFold 3 ^32^, which showed a conventional fungal laccase conformation with three core domains, domain I, G61–K183; domain-II, N184–M198 and Y245–G376; and domain-III, N377–E529, harboring four copper ions (Figure 5A-B). In addition, an auxiliary domain consisting of 42 amino acids (S199–M244) was observed (Figure 5A-B). The auxiliary domain region is located very close to the T1 copper, which is predicted to form the active site for PE binding, however its precise folding and interaction with the core domains could not be predicted. The T1 copper ion is located at the domain-III and tri-nuclear center consisting of one T2 copper and two T3 copper ions, located between domain-II and III. The axial ligands of the T1 copper site are reported to determine the redox potential of laccase enzymes, and *Pp*MmcO has an M505 residue as an axial ligand (Figure 5C). Laccases with methionine as the axial ligand generally have lower redox potentials than those with leucine or phenylalanine, and this reduced redox potential may compromise the oxidative capability of PpMmcO, thereby limiting its activity toward recalcitrant PE polymers ^33, 34^.

**Figure 5.**
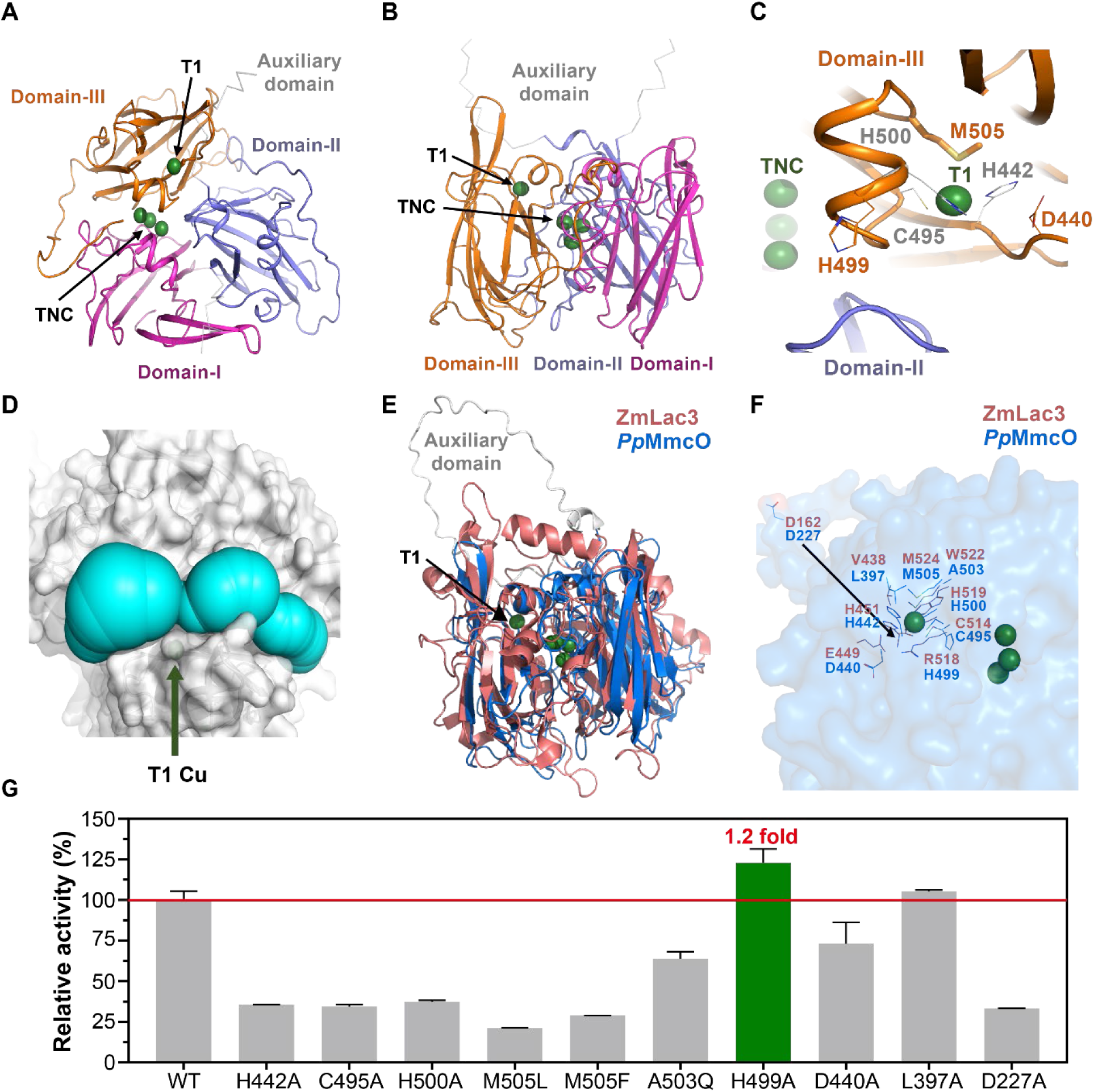
Predicted *Pp*MmcO^WT^ structure and comparative ABTS assay activities of its variants. The structure was predicted using AlphaFold 3. (**A and B**) The *Pp*MmcO^WT^ molecule is presented as cartoon diagram. Three core domains, domain-I, II, and III, are distinguished using magenta, blue, and orange, respectively. An auxiliary domain is presented as a grey ribbon. Four copper ions are drawn as green spheres with appropriate labels. (**C**) The T1 copper stabilization mode. The *Pp*MmcO^WT^ molecule is drawn as a cartoon diagram. Three core domains, domain-I, II, and III, are distinguished using magenta, blue, and orange, respectively. Four copper ions are drawn as green spheres with appropriate labels, and three core residues stabilizing the T1 copper ion are drawn as stick models with appropriate labels. The axial residue (M505) is shown as a stick model. (**D**) The putative substrate binding cavity in the *Pp*MmcO^WT^ structure. The predicted *Pp*MmcO^WT^ structure is shown as a grey surface model, while the predicted PE substrate binding cavities, calculated with CAVER software, are shown as cyan spheres. The T1 copper ion site, i.e., the catalytic site, is drawn as a labeled green sphere. (**E**) Structural alignment of the predicted *Pp*MmcO^WT^ model and the ZmLac3 (PDB ID: 6KLG) enzyme model. (**F**) The structure of residues forming *Pp*MmcO^WT^ superimposed with the residues of ZmLac3. (**G**) A comparison of the activities of the wild-type *Pp*MmcO and its variants based on the ABTS assay (*n* = 2, independent experiments).

Additionally, the CAVER software was used to predict the presence of a large substrate-binding cavity near the T1 copper site, which is the reported catalytic site of the laccase enzyme (Figure 5D) ^35^. The large substrate-binding cavity showed a flat conformation, potentially enabling it to bind hydrocarbon chains on a hydrophobic surface; however many hydrophilic residues, such as Q282, D371, E373, D441, K456, K474, and E477, are expected to create an unfavorable environment for the hydrophobic PE chains to easily bind to the active site (Figure 5D).

These structural insights led us to investigate how specific mutations in the amino acid residues near the T1 copper site might affect the enzymatic activity of *Pp*MmcO. The model structure of *Pp*MmcO was aligned with the structural model of ZmLac3 ^36^, a typical laccase (PDB: 6KLG), using SnapGene (GLS Biotech), producing an identity of 22.80% and a similarity of 38.78% (Figure S7). Both enzymes share the characteristic features of laccases, including one mononuclear copper (T1 Cu) site and one tri-nuclear center, and they preserve ten histidine residues and one cysteine residue that coordinate the four copper ions. Based on the alignment results, mutational analyses were performed to evaluate the effects of amino acid residue variants near the T1 copper site of *Pp*MmcO. The analyses targeted conserved residues (H442, C495, H500), the axial ligand (M505), T1 copper-surrounding residues (A503, H499, D440, L397), and the auxiliary domain (D227) (Figure 5E-F). Enzymatic activity was assessed using the ABTS assay (Figure 5G).

The C495A, H442A, and H500A variants involved in electron transfer near the T1 copper site showed a decrease in activity, achieving 21%, 35%, and 37% of the wild-type level, respectively. Similarly, the M505F and M505L variants, which were designed to achieve higher redox potential values, unexpectedly showed decreased activities, reaching 28% and 34%, respectively. The A503Q and D440A variants, predicted to form part of the substrate-binding pocket near the T1 copper, also exhibited reduced activities of 63% and 73%, respectively. Conversely, the L397A and H499A variants displayed slightly increased activity (105% and 122%, respectively) compared to the wild type, while the D227A variant, located within the auxiliary domain, displayed an activity decrease (33%). The D227 residue is predicted to play an important role in the enzyme’s function, although its exact structure could not be determined with confidence due to the low-confidence prediction in AlphaFold ^32^. These changes in activity demonstrate the intricate relationship between the structure and function of laccase enzymes and how modifications to amino acid residues near the T1 copper site can significantly impact enzymatic efficiency. However, because these insights are based on predicted structures, experimental high-resolution structural studies will be necessary to validate the folding of the auxiliary domain and the detailed configuration of the active site.

### 2.8. PELW powder, solution cast PELW film, and greenhouse PE film oxidation by the *Pp*MmcO H499A variant

Based on the ABTS assay, which confirmed that the H499A variant is expressed and catalytically active, we next evaluated whether this variant also exhibits oxidative activity toward PE substrates. Accordingly, reactions with PELW powder, solution-cast PELW film, and greenhouse PE film were conducted. SEM analysis of two film types, solution-cast PELW and greenhouse-cover PE films, treated with *Pp*MmcO^H499A^ revealed clear evidence of surface damage. The solution-cast PELW film treated with *Pp*MmcO^H499A^ (Figure 6A(b)) exhibited significant surface roughness and irregular textures compared to the smooth and unaltered surface of the control solution cast PELW film (Figure 6A(a)). Similarly, *Pp*MmcO^H499A^ treatment (Figure 6A(d)) resulted in observable damage and a roughened surface morphology in the greenhouse cover PE film, while the control greenhouse cover PE film maintained its smooth appearance (Figure 6A(c)). These results suggest that PpMmcO^H499A^, like *Pp*MmcO^WT^, can consistently induce structural modifications in solution cast PELW films and PE films.

**Figure 6.**
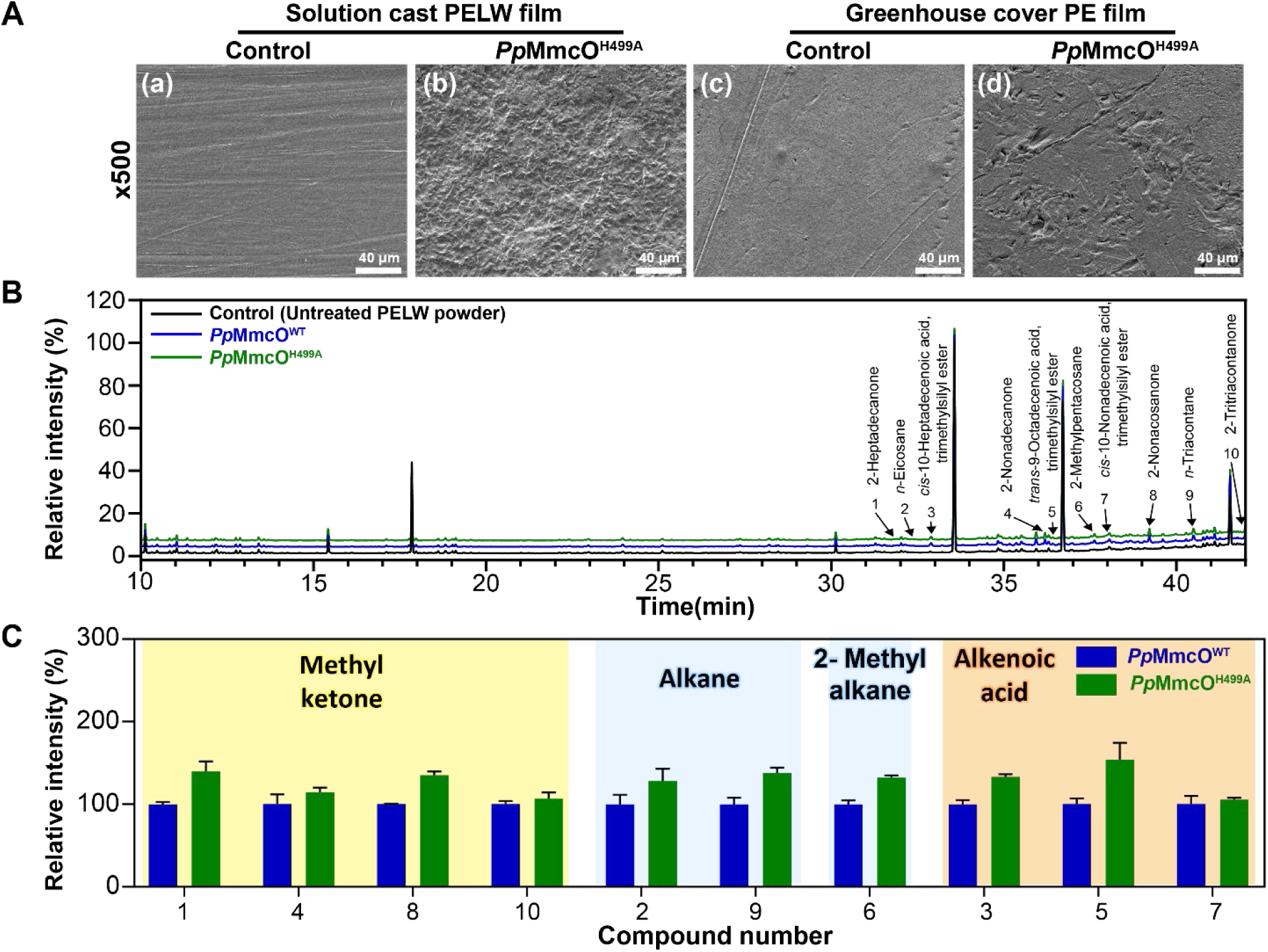
Degradation analysis of PELW powder, solution-cast PELW film, and greenhouse PE film by *Pp*MmcO^H499A^. (**A**) SEM images of (a and b) solution cast PELW and (c and d) greenhouse cover PE films treated with *Pp*MmcO^H499A^. (a and c) represent the untreated (control) PE film and (b and d) represent the *Pp*MmcO^H499A^-treated films. (**B**) GC-MS chromatograms of *Pp*MmcO^WT^ and *Pp*MmcO^H499A^ reaction solutions with PELW powder. Peaks are labeled with identified compounds. (**C**) Relative intensity (%) of releasing compounds after PELW degradation by comparing *Pp*MmcO^WT^ and *Pp*MmcO^H499A^. Compounds are grouped into methyl ketones, alkanes, 2-methyl alkane and alkenoic acids (*n* = 2, independent experiments).

The GC-MS analysis of the enzyme solution containing PELW powder and *Pp*MmcO^H499A^ showed higher intensities compare to that of *Pp*MmcO^WT^, indicating the presence of degradation products such as methyl ketones, alkanes, 2-methyl alkane, and alkenoic acids (Figure 6B, 6C and Table S5). This result suggests that *Pp*MmcO^H499A^ exhibits enhanced oxidative activity toward PE, as reflected by the higher intensities of released degradation products.

Compared with *Pp*MmcO^WT^, the *Pp*MmcO^H499A^ -treated PELW powder showed slightly increased hydrophilization, whereas boiled *Pp*MmcO^H499A^, like the untreated control, did not become suspended in the solution (Figure S8A).

Changes in the functional groups of PELW powder after reaction with *Pp*MmcO^H499A^ were analyzed using FT-IR. Both *Pp*MmcO^WT^ and *Pp*MmcO^H499A^ -treated PELW powders showed the presence of C−O stretching (carboxylic acid) and O−H stretching (hydroxyl group), as with the FT-IR results in Figure 3B. Slightly higher peak intensities were observed with *Pp*MmcO^H499A^ (Figure S8B).

To further confirm the activity of *Pp*MmcO^H499A^, enzymatic reactions were performed using PELW powder and the greenhouse PE film to assess weight loss. The PELW powder exhibited a weight loss of 5.0 ± 1.8% after treatment with *Pp*MmcO^H499A^, whereas the control (untreated PELW powder) showed only 0.5 ± 0.8%. In the case of the greenhouse PE film, the control sample did not show any weight loss, whereas the *Pp*MmcO^H499A^ -treated greenhouse PE film showed a mass-loss of 1.6 ± 0% (Figure S8C).

## 3. Discussion

In this study, we identified *P. polyethylenelyticus* JNU01 as a bacterium capable of growth on PELW and PE and sought to uncover the enzymes underlying this activity. A database-guided and transcriptome-based analysis led to the identification of *Pp*MmcO, a secreted multicopper oxidase (MmcO), as a gene notably upregulated under PELW conditions, highlighting the value of genome-wide approaches for discovering enzymatic activities on polymers. Among previous reports, two studies, those examining cytochrome P450 ^11^ and laccase-like multicopper oxidases ^15^, used the same 4 kDa PELW substrate from Sigma-Aldrich, which was also employed in this study. This shared substrate provides a useful basis for comparing enzymatic degradation mechanisms across studies.

While extracellular enzymes such as hydrolases and oxidases (e.g., laccases and peroxidases) have been proposed to contribute to PELW oxidation and depolymerization to small molecules, including dicarboxylic acids, alcohols, fatty acids, ketones, aldehydes, and esters ^37^, their precise mechanisms remain unresolved. Laccases, in particular, have been studied in the context of lignin degradation ^38^, a hydrophobic polymer with refractory C–C bonds, akin to linkages in polyolefins ^6, 39^. Some studies have suggested possible involvement of laccases in PE degradation ^13, 14, 40^, though key mechanistic details are lacking.

To elucidate the molecular mechanisms by which *Pp*MmcO contributes to PE oxidation, we propose a mechanism based on carbon radical oxidation reactions initiated by *Pp*MmcO (Figure 7), drawing inspiration from mechanistic studies from Gugumus ^41^. In the initial stage, *Pp*MmcO abstracts a hydrogen radical (H) from the polyethylene (PE) chain ^6, 42^, generating a carbon-centered radical (C·) (step 1), which reacts with molecular oxygen to form a peroxy radical (step 2). The peroxy radical then abstracts a hydrogen radical (H) from an adjacent carbon of PE, stabilizing as a hydroperoxide (step 3). Two possible pathways are then predicted for these hydroperoxide compounds. In the first pathway (steps 4–12), α,γ difunctional products are formed in PE ^41, 43–45^. These products consist primarily of dihydroperoxides (steps 4–6) and keto-hydroperoxides (steps 4–8). At the γ position, *Pp*MmcO abstracts a hydrogen radical (H·), generating a carbon radical (step 4). The hydroperoxide undergo further oxidation to form α,γ-dihydroperoxide structures (steps 5–6). The back-biting reaction at the α-carbon causes the radical to migrate along the PE chain (step 7), forming α,γ-keto-hydroperoxides (step 8), which further cyclize to form five-membered cyclic peroxides (step 9). Then, the five-membered cyclic peroxide compound cleaves into methyl ketone and carboxylic acid (step 10). The carboxylic acid undergoes further oxidation to produce alkenoic acid, again catalyzed by *Pp*MmcO (steps 11–12). In the second pathway (steps 13–18), the O−O bond in the hydroperoxide group breaks, forming a hydroxy radical (OH·) and another oxygen radical (step 13). The oxygen radical (O·) containing hydrocarbon chain then undergoes β-scission, resulting in methyl ketone and carbon radical containing hydrocarbon chain, which can be reduced by PEH to produce alkane (steps 14–15). Then, *Pp*MmcO abstracts a hydrogen radical (H) from the PE chain, generating a carbon radical (C ·). The carbon radical containing the hydrocarbon chain rearranges to produce a more stable structure, causing the radical position to shift (step 16) ^46^. During this process, a cyclopropane intermediate is formed (step 17). Then, cleavage of the C–C bond in the cyclopropane moiety may form 2-methyl alkane (step 18).

**Figure 7.**
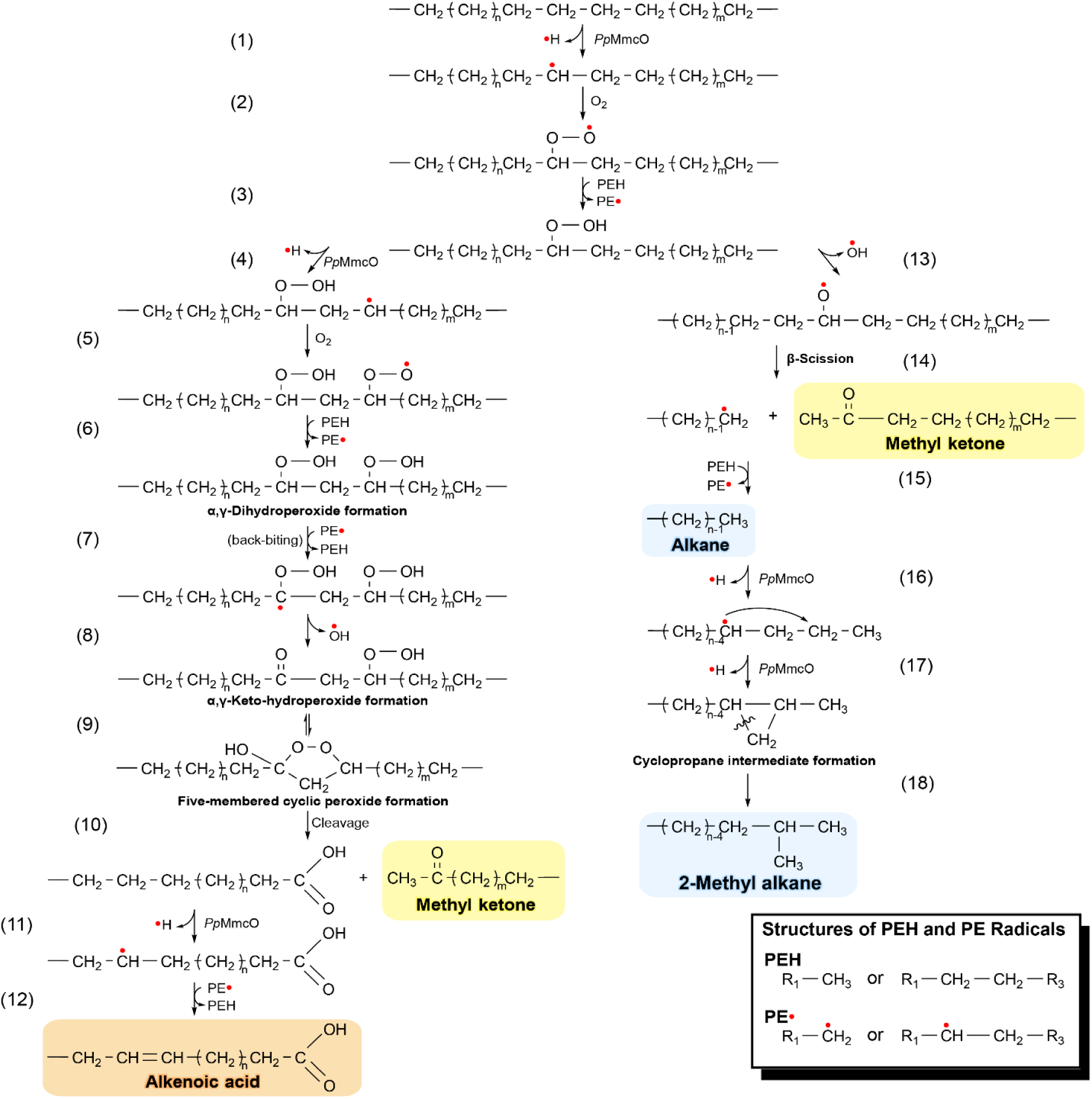
Proposed Mechanism of PE Degradation by *Pp*MmcO. The proposed mechanism for *Pp*MmcO catalyzed PE degradation involves two pathways initiated by carbon radical formation reaction.

This proposed dual-pathway mechanism, which remains to be evaluated in future work, suggests that *Pp*MmcO-catalyzed radical reactions can induce a cascading degradation of the PE chain. By enabling both in-chain hydroxylation and terminal oxidation, this proposed mechanism facilitates the formation of various degradation products, including alkenoic acids, methyl ketones, 2-methyl alkane, and alkanes, which were detected by GC-MS analysis of the enzyme solution after the *Pp*MmcO–PE reaction. These products are more accessible for biological processing, emphasizing that the radical-oxidization approach is a promising strategy for effective PE degradation and biological catabolism, similar to hybrid chemical and biological strategies proposed recently ^47–49^.

To gain more insight into the mechanism of PE degradation by *Pp*MmcO, we attempted to detect a five-membered cyclic peroxide, which is a proposed intermediate. The five-membered cyclic peroxides were detected (Figure S9) by bioluminescence based on the four-membered cyclic peroxide 1,2-dioxetane ^50^ detection method ^51, 52^. Accumulation of intermediates was evident with increasing concentration with increasing reaction time. This intermediate appears to play an important role in the production of various methyl ketones with different carbon chains which were observed in this study.

## 4. Conclusions

Collectively, these findings demonstrate that *P. polyethylenelyticus* JNU01 can utilize polyethylene (PE) as its sole carbon source, with its *Pp*MmcO initiating oxidative modification of PE. Specifically, a secreted multicopper oxidase, *Pp*MmcO, was shown to oxidatively transform polyethylene-like wax (PELW) in vitro, leading to the formation of oxygen-containing functional groups, the release of small-molecule oxidation products, and measurable weight loss of both PELW and post-use greenhouse PE films. These results support a model in which PpMmcO provides an oxidative entry point for the initial activation of the chemically inert C–C backbone of PE. Although several *Paenibacillus* species are known for their bioremediation potential (including the degradation of organic pollutants and recalcitrant compounds)^53–58^, the metabolic fate of the PE degradation products generated by *P. polyethylenelyticus* JNU01 remains unclear. Further investigation of the upregulated genes and physiological traits of this bacterium will be essential to determine whether it can assimilate or catabolize these intermediates, thereby contributing to a more complete understanding of microbial plastic degradation pathways.

## 5. Materials and Methods

### 5.1. Media, chemicals, PELW and PE preparation

Low-salt Luria–Bertani (LB) broth (10.0 g/L of tryptone, 5.0 g/L of yeast extract, and 5.0 g/L of NaCl) and M9 minimal broth (6.0 g/L of Na_2_HPO_4_, 3.0 g/L of KH_2_PO_4_, 0.50 g/L of NaCl, 1.0 g/L of NH_4_Cl, 240.7 mg/L of MgSO_4_, and 11.098 mg/L of CaCl_2_) with 0.1% (v/v) trace element solution (21.8 of mg/L CoCl_2_·6H_2_O, 21.6 mg/L of NiCl_2_·6H_2_O, 24.6 mg/L of CuSO_4_·5H_2_O, 1.62 g/L of FeCl_3_·6H_2_O, 0.78 g/L of CaCl_2_, and 14.7 mg/L of MnCl_2_·4H_2_O) were used.

A PE-like wax (PELW) substrate, marketed as PE, was purchased in powdered form from Sigma-Aldrich (Cat. No. 427772, St. Louis, MO, USA). This substrate had an average M_w_ of ∼4,000 and an average M_n_ of ∼1,700, as measured using gel permeation chromatography, and is referred to throughout as PELW. The preparation of the PELW polymer model substrate for powder and films was carried out using the method reported in our previous study ^11^. Briefly, PELW powder was re-precipitated by dissolving it in chloroform at 55 °C and precipitating it in methanol, followed by vacuum drying. PELW films were fabricated by dissolving the re-precipitated PELW in heptane at 70 °C and casting the solution into Teflon dishes for solvent evaporation. In addition, the post-use greenhouse cover PE film used in this study was a PE film (Ilsin Chemical Industry, Ansan, Republic of Korea) with an Mw of 119,000 and an Mn of 18,000, as determined by gel permeation chromatography (GPC) at the Korea Polymer Testing & Research Institute (KOPTRI, Seoul, Republic of Korea). 2,2’-Azino-bis(3-ethylbenzothiazoline-6-sulfonic acid) (ABTS) was also purchased from Sigma-Aldrich, ethanol (99.9 %) was purchased from Thermo Fisher Scientific (Waltham, MA USA), and BSTFA-TMCS (99:1) [Derivatizing Reagent for GC] was purchased from TCI chemicals (Tokyo, Japan). Chloroform (CHCl_3_, 99%), methanol (≥ 99.8%), heptane (≥ 98%), and Ethyl acetate (99.5%) were purchased from Duksan (Ansan, Republic of Korea).

### 5.2. Screening and identification of PELW-degrading microorganism

Environmental samples (soil and waste) were obtained from the metropolitan sanitary landfill located in Gwangju, Republic of Korea. Soil and waste buried approximately 20 meters underground were obtained using a drilling rig. The screening of PELW-degrading microorganisms was performed using the method described in our previous study ^11^. PELW-degrading microorganisms were screened by culturing landfill-derived soil samples in M9 minimal medium containing 1 g/L PELW as the sole carbon source. Colonies showing growth were re-streaked, and selected isolates were identified by 16S rRNA sequencing. The growth of colonies was monitored over 40 days in PELW-containing liquid medium.

The fold change of the OD_600_ value of strain 2GT37 (*Paenibacillus* sp. JNU01) was measured using a VICTOR Nivo^TM^ Multimode Microplate Reader (PerkinElmer, Waltham, MA, USA) every 10 days. Afterwards, genomic DNA was extracted from strain 2GT37 using a HiGene^TM^ Genomic DNA Prep Kit (Biofact, Daejeon, Republic of Korea), and 16s rRNA sequencing was performed at Solgent (Daejeon, Republic of Korea). Additionally, a phylogenetic tree was obtained using MEGA X (v11.0.13) software. Genomic DNA extraction, 16S rRNA sequencing, and phylogenetic tree analysis were conducted as outlined in our previous study ^11^.

### 5.3. Cultivation of PELW-degrading microorganism and media analysis

*Paenibacillus* sp. JNU01 was inoculated into LB medium and cultured overnight at 28 °C. The culture was then centrifuged at 3,500 g for 20 min at 4 °C, and the cells were washed using M9 minimal medium. The washed *Paenibacillus* sp. JNU01 cells were then resuspended in M9 minimal medium, adjusting the initial OD_600_ to 0.20. The growth curve of *Paenibacillus* sp. JNU01 was monitored in cultures with the powder of the PELW model substrate at concentrations of 10, 30, and 50 mg/mL. Cultivation was carried out in 125-mL flasks containing 30 mL of M9 minimal medium at 28 °C and 200 rpm in a shaking incubator for 30 days. For PELW films, the same medium was used with an initial OD_600_ of 2.0, and cultures were performed in 50-mL flasks containing 10 mL of M9-TES medium, incubated at 28 °C and 120 rpm in a shaking incubator for 30 days. The OD_600_ value measured every 5 days using a UV-Vis spectrophotometer (UV-1900, Shimadzu, Kyoto, Japan). Dissolved oxygen concentrations in M9-TES medium were monitored using a biochemical oxygen demand probe and a DO meter (YSI 4100 BOD IDS PROBE & YSI 4010-1W DO meter, respectively, YSI Incorporated, Yellow Springs, Ohio, USA), measuring every 5 days for 10 days.

Additionally, GC-MS analyses were performed on the culture media during growth on the PELW model substrate. The media was first extracted with an equal volume of ethyl acetate (EA). The EA layer was then concentrated 50-fold, filtered, and analyzed using GC-MS (Shimadzu-QP2020 NX, Shimadzu) with an electron impact ionization source and a DB-5MS capillary column (Agilent, Santa Clara, CA, USA). The column temperature was maintained at 40 °C for 3 min, increased from 40 °C to 280 °C at a heating rate of 6 °C min^−1^ and maintained at 280 °C for 4 min. The injector and detector were held at 300 °C and 260 °C, respectively. The chemical structures of the products were library-matched using the NIST/WILEY database.

### 5.4. Characterization of chemical and physical changes to PELW powder, solution cast PELW film, and greenhouse cover PE film after treatment with *Paenibacillus* sp. JNU01

The three forms of PE (PELW powder, solution-cast PELW film, and greenhouse cover PE film) were each incubated with *Paenibacillus* sp. JNU01 in M9 minimal medium for 30 days. The powder form of the PELW model substrate was filtered using Whatman^TM^ Grade 1 Qualitative Filter Paper (90 mm, CAT No. 1001-090, GE Healthcare Life Science, Maidstone, UK). The filtered powder was sequentially washed with methanol (four times), 1 M NaOH (five times), and distilled water (five times), followed by vacuum drying. To identify chemical functional groups, Fourier Transform Infrared Spectroscopy (FT-IR) spectroscopy was used, measuring wavenumbers in the range of 4,000–500 cm^−1^ on a Spectrum 100 instrument (PerkinElmer).

Solution cast-PELW films and greenhouse cover PE films were washed in a 2% SDS solution for 4 h, thoroughly washed four times with methanol, and then vacuum-dried. To create a thin conductive layer on the solution cast-PELW film and greenhouse-cover PE film, coating was performed using platinum at 10 mA for 60 s. Then, changes in the surface of the solution-cast PELW films and greenhouse cover PE films before and after treatment with the strain were observed using field-emission scanning electron microscope (FE-SEM) (JSM-7900F, JEOL, Tokyo, Japan) with an accelerating voltage of 2.0 kV at the Engineering Practice Education Center (Chonnam National University). In addition, surface hydrophilicity (or hydrophobicity) was measured using water contact angle (WCA) analysis equipment (Phoenix 300, Seo, Suwon, Republic of Korea) to assess changes due to surface roughness and chemical structure alterations.

### 5.5. Genome sequencing and analysis of *P*. *polyethylenelyticus* JNU01

To obtain the genome of the PELW-degrading bacterium *P. polyethylenelyticus* JNU01, cells were cultured overnight in LB medium at 28 °C. Genomic DNA was extracted using a HiGene™ Genomic DNA Prep Kit (Biofact), and its quality and concentration were assessed using a NanoDrop spectrophotometer (Thermo Fisher Scientific) and Quant-IT PicoGreen (Invitrogen, CA, USA). A 20 kb PacBio SMRTbell library was prepared, with size distributions confirmed using a Bioanalyzer 2100 (Agilent). High-molecular-weight DNA was sheared and purified for optimal library insert size using g-TUBEs (Covaris Inc., MA, USA) and AMPure PB magnetic beads (Beckman Coulter Inc., CA, USA). The library was prepared using a PacBio DNA Template Prep kit 1.0, and sequencing was performed on a PacBio RS II platform (Pacific Bioscience, CA, USA) with C4 chemistry, capturing 240-minute movies for each SMRT cell, by Macrogen (Seoul, Republic of Korea). A single SMRT cell generated 134,300 subreads, yielding 1.31 Gb of data with an average length of 9,787 bp and an N50 length of 14,569 bp. For high-quality sequencing, a paired-end DNA library with an average insert size of 350 bp was prepared using Covaris shearing and an Illumina TruSeq Nano DNA library preparation kit. Sequencing on an Illumina HiseqX platform (Macrogen) produced approximately 4.9 million 2 × 151 bp paired-read sequences. Adapter sequences were trimmed and low-quality reads were removed using Hierarchical Genome Assembly Process 3 (HGAP3 v3.0)^59^ and Trimmomatic (v0.36)^60^ with the default settings. The resulting PacBio SMRT sequencing reads were *de novo* assembled using HGAP3 with pre-assembly (minimum seed read length = 6,000 bp, minimum polymerase read quality = 0.80, minimum polymerase read length = 100 bp). The PacBio assembled genome was polished using Pilon v1.21^61^, with the default parameters, and the trimmed high-quality Illumina reads (1.02 Gb of data providing approximately 148× coverage of the estimated genome size). The final assembled genome was annotated using the rapid prokaryotic genome annotation tool Prokka (v1.14.5) ^62^. The circular genome was 6,876,188 bp in length and consisted of 6,113 predicted genes, including 5,973 protein-coding genes, 36 rRNA genes, and 103 tRNA genes. Circular genome visualization was performed using Proksee ^63^.

### 5.6. Transcriptome sequencing and analysis of *P. polyethylenelyticus* JNU01

After culturing overnight in LB medium at 28 °C and 200 rpm in a shaking incubator, *P. polyethylenelyticus* JNU01 cells were centrifuged for 20 min at 3,500 g, washed with M9 minimal broth, and inoculated into M9 minimal broth media supplemented with either 0.4% (v/v) glucose, for the control group, or 30 mg/mL of model PELW powder, for the experimental group. Both groups had an initial OD_600_ of 0.20 and were cultured to the mid-exponential phase at 28 °C and 200 rpm in a shaking incubator. Total RNA was extracted using the Monarch® Total RNA Miniprep Kit (New England BioLabs, Ipswich, MA, USA) and quality-checked using a Bioanalyzer 2100 system (Agilent Technologies) and Nanodrop ND-2000 Spectrophotometer (Thermo Scientific). The rRNA was then removed using the NEB Next rRNA Depletion Kit (New England BioLabs) and RNA libraries were prepared using the TruSeq Stranded Total RNA Library Prep Kit with Ribo-Zero Gold (Illumina, Inc., USA) and screened for quality and quantity using TapeStation HS D1000 Screen Tape (Agilent Technologies) and a StepOne Real-Time PCR System (Life Technologies, Inc., USA). High-throughput sequencing was performed as paired-end 151 bp sequencing on a NovaSeq 6000 system (Illumina, Inc.).

Raw sequence reads were preprocessed using Trimmomatic (v0.39) ^60^ to remove low-quality bases and adapter sequences. To improve the read alignment efficiency, a genome index was constructed for *P. polyethylenelyticus* JNU01 based on the *de novo* assembled genomic data using the “bowtie2-build” command in Bowtie2 (v2.2.1) ^64^. Cleaned RNA-Seq paired-end reads were then aligned to this genome using Bowtie2 with the default settings. The resulting binary alignment map (BAM) files were processed with StringTie (v2.2.1) and prepDE ^65^. Differentially-expressed genes were identified using the edgeR package (v3.38.4) ^66^, applying a minimum read count of 10in at least one sample and a false discovery rate (*q*-value) < 0.05. Transcript abundance was quantified using either transcripts per kilobase million or counts per million mapped reads.

### 5.7. Bioinformatics analysis

Potentially relevant oxidative enzymes were identified through analysis of known enzymes from peer-reviewed literature and curated databases, including PlasticDB ^26^, the plastics microbial biodegradation database (PMBD) ^27^, and the plastic-active enzymes database (PAZY) ^28^. The amino acid sequences of these enzymes were retrieved from the UniProtKB/Swiss-Prot database using their EC numbers. OrthoMCL (v2.0.9) ^67^ was then used to identify corresponding orthologous genes in *P. polyethylenelyticus* JNU01. Putative secreted proteins, presumed to be potentially involved in PE-degradation, were predicted using SignalP (v6.0) ^31^.

Multicopper oxidase sequences were retrieved from the UniProt/SwissProt database using EC numbers for laccases (1.10.3.2), ascorbate oxidases (1.10.3.3), and ferroxidases (1.16.3.1). After filtering out amino acid sequences shorter than 100 and longer than 1,000, 83 laccase, 6 ascorbate oxidase, and 109 ferroxidase sequences were retained. To investigate phylogenetic relationships, three previously reported polyethylene-degrading multicopper oxidases were included (Figure S4)^15, 16^. Sequences were aligned using MAFFT (v7.520) using the “—auto” parameter ^68^, followed by quality checks and format conversion using trimAl (v1.2rev59) with the “-gt” parameter set to 0.3 ^69^. ModelFinder ^70^ was used to determine the best-fit model before reconstructing the maximum-likelihood (ML) tree. The ML tree was inferred using IQ-TREE (v2.3.4) with 1,000 bootstrap resamples, utilizing the Q.pfam+R6 model, as selected by the Bayesian information criterion ^71^.

### 5.8. Gene cloning and the site-directed mutagenesis of *Pp*MmcO

The nucleotide sequence corresponding to the signal peptide in the DNA sequence encoding *Pp*MmcO was predicted and removed using SignalP (v6.0) ^31^. The gene encoding the signal peptide-removed *Pp*MmcO (1,530 bp) was amplified from the genomic DNA from *P. polyethylenelyticus* JNU01 using PrimeSTAR® Max DNA Polymerase Ver.2 (TaKara, Kusatsu, Shiga, Japan). Amplified DNA fragments were purified using the Wizard® SV Gel and PCR Clean-Up System (Promega, Madison, WI, USA), and then the *Pp*MmcO gene and pProEX HTA vector were ligated into the plasmid using Gibson Assembly® Master Mix (New England Biolabs, Ipswich, MA, USA). The resulting plasmid was transformed into *E. coli* strain C2566 in an electroporator (MicroPulser, Bio-Rad, Hercules, CA, USA) and spread on LB agar containing 50 μg/mL of ampicillin. The sequence of the constructed plasmid was confirmed by Sanger sequencing (Macrogen). Site-directed mutagenesis of *Pp*MmcO was performed using Muta-Direct™ Site Directed Mutagenesis kits (iNtRON Biotechnology, Seongnam, Republic of Korea). The transformation and sequencing of the *Pp*MmcO variants were performed using the same methods. The primer sequences used for cloning and site-directed mutagenesis are listed in Table S2.

### 5.9. Overexpression and purification of wild-type *Pp*MmcO and its variants

Wild-type and variant *Pp*MmcO-transformed *E. coli* C2566 were cultured overnight in LB media at 37 °C and 200 rpm in a shaking incubator. Transformed *E. coli* C2566 were inoculated at 1% (v/v) into 500 mL of LB media containing 50 μg/mL of ampicillin and cultured in a shaking incubator at 37 °C and 200 rpm until the OD_600_ reached 0.6–0.8. Transformed *E. coli* C2566 were then induced by the addition of 0.5 mM ITPG (isopropyl-β-D-thiogalactopyranoside) supplemented at 1.0 mM with CuSO_4_·5H_2_O, and the culture was incubated in a shaking incubator at 37 °C and 200 rpm for 3 hours followed by 20 °C and 150 rpm for about 16 hours. *E. coli* C2566 overexpressing recombinant *Pp*MmcO was harvested by centrifugation at 4 °C and 3,500 × g for 20 min. The pellet was resuspended in a 5 mM imidazole, 40 mM Tris-HCl buffer (pH 7.5), and disrupted by ultra-sonication (Fisher Scientific). After centrifugation at 4 °C and 12,000 g, the crude extract was prepared, loaded onto a WorkBeads^TM^ 40 Ni-NTA column (Bio-Works, Uppsala, Sweden), and eluted with a 300 mM imidazole, 40 mM Tris-HCl buffer (pH 7.5). The eluted solution was centrifuged at 12,000 × g for 10 min at 4 °C, and the supernatant was applied to a HiTrap Q HP anion exchange chromatography column (Cytiva, Marlborough, MA, USA) and equilibrated with buffer A (40 mM Tris-HCl at pH 7.5) using an NGC Fast Protein Liquid Chromatography (FPLC) system (Bio-Rad, Hercules, CA, USA). The bound protein was eluted with buffer B (40 mM Tris-HCl and 2 M NaCl at pH 7.5) at a flow rate of 2 mL min^−1^. The eluted protein was then placed in an ultracentrifugation filtration tube (50 kDa molecular weight cutoff, Amicon, Sigma-Aldrich, St. Louis, MO, USA) and dialyzed against 50 mM potassium phosphate buffer (pH 7.5). The molar mass of the purified enzyme was determined by SDS-PAGE.

### 5.10. Wild-type and variant *Pp*MmcO activity analyses

The wild-type and variant *Pp*MmcO activities were evaluated by monitoring the change in absorbance at 420 nm over 2 min using ABTS as a substrate and a UV-Vis spectrophotometer (UV-1900, Shimadzu) ^72^. One unit of enzyme activity was defined as the amount of enzyme that oxidized 1.0 μmol of ABTS substrate per min^73^. Enzyme activity was calculated using a slight modified version of the formula adopted from Leonowicz and Grzywnowicz ^74^:

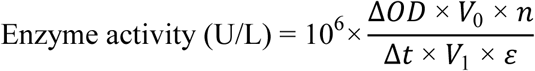

where Δ*OD* is the change in absorbance, *V*_0_ is the total enzyme reaction volume (mL), *n* is dilution ratio of the enzyme solution, Δ*t* is the time (min), *V*_1_ is the volume of enzyme (mL), and ε is the molar extinction coefficient (ε = 36,000 M^−1^ cm^−1^).

### 5.11. Effects of temperature, pH, and copper ion concentration on the activity and thermal stability of the wild-type *Pp*MmcO

The effect of temperature on *Pp*MmcO^WT^ activity was monitored at various temperatures (25-80°C in 5 °C increments) in 1.0 mL of 20 mM Na-acetate buffer (pH 4.0) containing CuSO_4_·5H_2_O at 2.0 mM, ABTS at 0.50 mM, and 0.010 mg/mL of enzyme. The enzyme solutions, excluding ABTS, were incubated in water baths at various temperatures for 30 min before initiating the reaction by adding ABTS. Next, to evaluate the effect of pH on *Pp*MmcO^WT^ activity, we used 20 mM Glycine-HCl buffer (pH 2.5–3.6) or 20 mM Na-acetate buffer (pH 3.6–5.0), with pH values collectively ranging from 2.5 to 5.0. The reaction mixture, which included the appropriate buffer, along with CuSO_4_·5H_2_O at 2.0 mM, ABTS at 0.5 mM, and 0.010 mg/mL of enzyme in a total reaction volume of volume of 1.0 mL. The enzyme activities of reaction solutions, excluding ABTS, were measured by adding ABTS after incubation in a water bath at 65°C for 30 min. To investigate the effect of copper ion concentration on *Pp*MmcO^WT^ enzyme activity, the enzyme activity was monitored in 1.0 mL of a 0.50 mM ABTS, 20 mM Na-acetate buffer (pH 3.6) containing CuSO_4_·5H_2_O at various concentrations (up to 10 mM) and 0.010 mg/mL of enzyme. The enzyme activities of reaction solutions, excluding ABTS, were measured by adding ABTS after incubation in a water bath at 65 °C for 30 min. The long-term thermal stability of *Pp*MmcO^WT^ was monitored by measuring the residual activity at three different temperature conditions (30 °C, 50 °C, and 65 °C) after incubating 1.0 mL of enzyme mixture (20 mM Na-acetate buffer (pH 3.6), 2.0 mM CuSO_4_, and 0.010 mg/mL of enzyme) without ABTS for 1, 2, 4, 6, 16, and 24 h, and every 24 h after up to 120 h. The enzyme activities of reaction solutions, excluding ABTS, were measured by adding ABTS after incubation in a water bath at 65 °C for 30 min.

### 5.12. Oxidative activities of wild-type and variant *Pp*MmcO on ABTS, PELW powder, solution-cast PE film, and greenhouse-cover PE film

The enzyme activities of the *Pp*MmcO^WT^ and its variants for ABTS assay were monitored by measuring 1.0 mL of enzyme solutions (20 mM Na-acetate buffer (pH 3.6), 2.0 mM CuSO_4_, 0.50 mM ABTS, and 0.010 mg/mL of enzyme). The reaction solution, excluding ABTS, was first incubated in a water bath at 65 °C for 30 min, after which ABTS was added to initiate the reaction. Enzyme activity was monitored by measuring the absorbance at 420 nm over 2 min using a UV-Vis spectrophotometer (UV-1900, Shimadzu).

The oxidative activities of *Pp*MmcO^WT^ and its variant, *Pp*MmcO^H499A^, on PELW was analyzed using three forms of PE: PELW powder (10 mg/mL), solution cast PELW film (0.5 cm × 0.5 cm) and greenhouse cover PE film (0.5 cm × 0.5 cm). Each PE substrate was mixed into 2 mL of reaction buffer consisting of 2 mM CuSO_4_ and 20 mM Na-acetate (pH 3.6) containing one of the purified enzymes at specific concentrations. The reactions were generally performed at an enzyme concentration of 0.20 mg/mL, although in some cases, *Pp*MmcO^WT^ was tested at 0.20, 1.0, and 3.0 mg/mL. The enzyme–PE reaction solutions were incubated at 50 °C in a water bath for 24 hours. As negative controls, heat-inactivated *Pp*MmcO^WT^ and *Pp*MmcO^H499A^ (0.20 mg/mL each, boiled at 95 °C for 10 min) were also included in the PELW powder reactions.

After the reaction, the washing procedures were performed separately depending on the three types of PE substrate. For PELW powder, samples were washed sequentially with methanol (4 times), 1 M NaOH (5 times), and distilled water (5 times), followed by vacuum drying. For solution-cast PELW films and greenhouse-cover PE films, samples were washed in a 2% SDS solution for 4 h, thoroughly rinsed four times with methanol, and then vacuum-dried. The washed and dried samples were then analyzed for chemical and physical changes using FT-IR, SEM, and WCA, as described in the Methods subsection “Characterization of chemical and physical changes to PELW powder, solution cast PELW film, and greenhouse cover PE film after treatment with *Paenibacillus* sp. JNU01” Weight loss was measured by comparing the combined mass of the test tube and PELW powder or greenhouse PE film before and after enzymatic reactions with *Pp*MmcO^WT^ and *Pp*MmcO^H499A^, using a precision analytical balance (Pioneer^TM^ PX124KR, Ohaus, Parsippany, NJ, USA). For GC-MS analyses, the enzyme reaction solutions were extracted with an equal volume of EA, and the EA layer was concentrated 30-fold and derivatized with BSTFA-TMCS at 55 °C for 40 min. The derivatized samples were then analyzed using GC-MS as described in the Methods subsection “Cultivation of PELW-degrading microorganism and media analysis”.

### 5.13. Luminescence measurement of wild-type *Pp*MmcO

The enzyme solution (3 mL) contained 20 mM Na-acetate buffer (pH 3.6), 2 mM CuSO_4_, and 0.20 mg/mL *Pp*MmcO^WT^, and PELW powder at a concentration of 10 mg/mL. The enzyme solution was incubated in a 50 °C water bath. Luminescence intensity was measured at 450 nm using a VICTOR Nivo^TM^ Multimode Microplate Reader (PerkinElmer) ^51, 52^. Samples were collected and analyzed at 5, 10, 20, and 30 min after the initiation of the reaction. The control solution containing PELW powder without the enzyme was used.

## ASSOCIATED CONTENT

### Supporting information

Supplementary Figures include: identification and phylogenetic analysis of PELW-biodegrading strains (Fig. S1); evaluation of extended washing procedures on FT-IR spectra of PELW powders treated with *Paenibacillus polyethylenelyticus* JNU01 (Fig. S2); transcriptomic heatmap of candidate genes potentially involved in PELW biodegradation (Fig. S3); phylogenetic analysis of *Pp*MmcO among multicopper oxidases (Fig. S4); SDS-PAGE analysis of *Pp*MmcO wild type and variants (Fig. S5); optimization of reaction conditions for *Pp*MmcO activity, including temperature, pH, copper concentration, and thermal stability (Fig. S6); amino acid sequence alignment of ZmLac3 and *Pp*MmcO (Fig. S7); chemical and physical changes in PELW powder and PE films following *Pp*MmcO treatment, including FT-IR spectra and weight-loss analysis (Fig. S8); and luminescence measurements associated with *Pp*MmcO activity in the presence of PELW powder (Fig. S9).

Supplementary Tables include: GC–MS-based identification of metabolites detected from *Paenibacillus* sp. JNU01 cultured in PELW-containing media (Table S1); primer sequences used for cloning and site-directed mutagenesis of *Pp*MmcO and its variants (Table S2); amino acid sequences of *Pp*MmcO wild type and variants with signal peptides and mutation sites annotated (Table S3); GC–MS analysis of compounds generated by the *Pp*MmcO wild type in enzyme reaction solutions containing PELW powder at different enzyme concentrations (Table S4); and comparative GC–MS analysis of reaction products generated by the *Pp*MmcO wild type and the H499A variant (Table S5). Supplementary Data include: average nucleotide identity (ANI) analysis among related species and strains (Supplementary Data S1); RNA-seq analysis statistics (Supplementary Data S2); lists of upregulated (Supplementary Data S3) and downregulated (Supplementary Data S4) differentially expressed genes in response to PELW; and amino acid sequences of enzymes used for phylogenetic analysis of *Pp*MmcO (Supplementary Data S5).

## AUTHOR INFORMATION

### Author Contributions

**Seung-Do Yun:** Conceptualization, Methodology, Software, Formal analysis, Investigation, Visualization, Writing – original draft. **Seongmin Kim:** Methodology, Software, Formal analysis, Investigation, Visualization, Writing – original draft. **Seong Jin An:** Methodology, Investigation, Visualization, Writing – original draft. **Hyun-Woo Kim:** Investigation, Data curation. **Jin-Hee Cho:** Investigation, Data curation. **Hyeoncheol Francis Son:** Formal analysis, Investigation, Writing – original draft. **Chul-Ho Yun:** Investigation, Methodology, Writing – review & editing. **Bong Hyun Sung:** Investigation, Funding acquisition. **Gregg T. Beckham**: Methodology, Writing – review & editing. **Won Seok Chi:** Methodology, Supervision, Writing – review & editing. **Chungoo Park:** Methodology, Supervision, Writing – review & editing. **Soo-Jin Yeom:** Conceptualization, Methodology, Funding acquisition, Project administration, Supervision, Writing – review & editing. †These authors contributed equally to this work

### Notes

The authors declare no competing interests.

## ACKNOWLEDGMENT

**Funding Sources:** This research was supported by the Bio & Medical Technology Development Program (RS-2024-00440681), Enzyme Engineering for Next Generation Biorefinery Program (NRF-2022M3J5A1056169 and NRF-2022M3J5A1085239), Synthetic Biology Core Technology Development Program (RS-2024-00398252), NRF grant (RS-2023-00208002) and Korea Basic Science Institute (National Research Facilities and Equipment Center) grant (RS-2024-00398582) funded by the Korean government (MSIT). Funding to GTB was provided as part of the Bio-Optimized Technologies to keep Thermoplastics out of Landfills and the Environment (BOTTLE) Consortium supported by the Advanced Materials and Manufacturing Technologies Office (AMMTO) and Bioenergy Technologies Office (BETO) under contract no. DE-AC36-08GO28308 with the National Renewable Energy Laboratory (NREL). The views expressed in the article do not necessarily represent the views of the DOE or the U.S. Government. We thank Gustav Vaaje-Kolstad for helpful discussions regarding the nature of the 4 kDa PELW substrate from Sigma-Aldrich.

## REFERENCES

(1) Vidal, F.; van der Marel, E. R.; Kerr, R. W. F.; McElroy, C.; Schroeder, N.; Mitchell, C.; Rosetto, G.; Chen, T. T. D.; Bailey, R. M.; Hepburn, C.; Redgwell, C.; Williams, C. K., Designing a circular carbon and plastics economy for a sustainable future. Nature. 2024, 626 (7997), 45–57.

(2) Ellis, L. D.; Rorrer, N. A.; Sullivan, K. P.; Otto, M.; McGeehan, J. E.; Román-Leshkov, Y.; Wierckx, N.; Beckham, G. T., Chemical and biological catalysis for plastics recycling and upcycling. Nat. Catal. 2021, 4 (7), 539–556.

(3) Stubbins, A.; Law, K. L.; Muñoz, S. E.; Bianchi, T. S.; Zhu, L., Plastics in the Earth system. Science. 2021, 373 (6550), 51–55.

(4) Borrelle, S. B.; Ringma, J.; Law, K. L.; Monnahan, C. C.; Lebreton, L.; McGivern, A.; Murphy, E.; Jambeck, J.; Leonard, G. H.; Hilleary, M. A.; Eriksen, M.; Possingham, H. P.; De Frond, H.; Gerber, L. R.; Polidoro, B.; Tahir, A.; Bernard, M.; Mallos, N.; Barnes, M.; Rochman, C. M., Predicted growth in plastic waste exceeds efforts to mitigate plastic pollution. Science. 2020, 369 (6510), 1515–1518.

(5) Chamas, A.; Moon, H.; Zheng, J.; Qiu, Y.; Tabassum, T.; Jang, J. H.; Abu-Omar, M.; Scott, S. L.; Suh, S., Degradation Rates of Plastics in the Environment. ACS Sustain. Chem. Eng. 2020, 8 (9), 3494–3511.

(6) Liu, L.; Zhang, M.; Xu, H.; Blažević, Z. F.; Ballerstedt, H.; Blank, L. M., Challenges of polyethylene (PE) biodegradation – A perspective. Biotechnol. Adv. 2025, 85, 108717.

(7) Roy, P. K.; Hakkarainen, M.; Varma, I. K.; Albertsson, A.-C., Degradable polyethylene: fantasy or reality. Environ. Sci. Technol. 2011, 45 (10), 4217–4227.

(8) Yeom, S.-J.; Le, T.-K.; Yun, C.-H., P450-driven plastic-degrading synthetic bacteria. Trends Biotechnol. 2022, 40 (2), 166–179.

(9) Albertsson, A.-C.; Andersson, S. O.; Karlsson, S., The mechanism of biodegradation of polyethylene. Polym. Degrad. Stab. 1987, 18 (1), 73–87.

(10) Yoon, M. G.; Jeon, H. J.; Kim, M. N., Biodegradation of polyethylene by a soil bacterium and AlkB cloned recombinant cell. J. Biorem. Biodegrad. 2012, 3 (4), 1–8.

(11) Yun, S.-D.; Lee, C. O.; Kim, H.-W.; An, S. J.; Kim, S.; Seo, M.-J.; Park, C.; Yun, C.-H.; Chi, W. S.; Yeom, S.-J., Exploring a New Biocatalyst from *Bacillus thuringiensis* JNU01 for Polyethylene Biodegradation. Environ. Sci. Technol. Lett. 2023, 10 (6), 485–492.

(12) Son, J.-S.; Lee, S.; Hwang, S.; Jeong, J.; Jang, S.; Gong, J.; Choi, J. Y.; Je, Y. H.; Ryu, C.-M., Enzymatic oxidation of polyethylene by *Galleria mellonella* intestinal cytochrome P450s. J. Hazard. Mater. 2024, 480, 136264.

(13) Gao, R.; Liu, R.; Sun, C., A marine fungus *Alternaria alternata* FB1 efficiently degrades polyethylene. J. Hazard. Mater. 2022, 431, 128617.

(14) Zhang, A.; Hou, Y.; Wang, Q.; Wang, Y., Characteristics and polyethylene biodegradation function of a novel cold-adapted bacterial laccase from Antarctic sea ice psychrophile *Psychrobacter* sp. NJ228. J. Hazard. Mater. 2022, 439, 129656.

(15) Zampolli, J.; Mangiagalli, M.; Vezzini, D.; Lasagni, M.; Ami, D.; Natalello, A.; Arrigoni, F.; Bertini, L.; Lotti, M.; Di Gennaro, P., Oxidative degradation of polyethylene by two novel laccase-like multicopper oxidases from *Rhodococcus opacus* R7. *Environ*. Technol. Innov. 2023, 32, 103273.

(16) Zhang, X.; Feng, X.; Lin, Y.; Gou, H.; Zhang, Y.; Yang, L., Degradation of polyethylene by *Klebsiella pneumoniae* Mk-1 isolated from soil. Ecotoxicol. Environ. Saf. 2023, 258, 114965.

(17) Jendrossek, D., Polyethylene and related hydrocarbon polymers (“plastics”) are not biodegradable. N. Biotechnol. 2024, 83, 231–238.

(18) Kim, H.-W.; Jo, J. H.; Kim, Y.-B.; Le, T.-K.; Cho, C.-W.; Yun, C.-H.; Chi, W. S.; Yeom, S.-J., Biodegradation of polystyrene by bacteria from the soil in common environments. J. Hazard. Mater. 2021, 416, 126239.

(19) Stepnov, A. A.; Lopez-Tavera, E.; Klauer, R.; Lincoln, C. L.; Chowreddy, R. R.; Beckham, G. T.; Eijsink, V. G. H.; Solomon, K.; Blenner, M.; Vaaje-Kolstad, G., Revisiting the activity of two poly(vinyl chloride)- and polyethylene-degrading enzymes. Nat. Commun. 2024, 15 (1), 8501.

(20) Devi, P.; Fatma, S.; Parveen, H.; Bishnoi, A.; Singh, R., Spectroscopic analysis, first order hyperpolarizability, NBO, HOMO and LUMO analysis of 5-oxo-1-phenylpyrrolidine-3-carboxylic acid: Experimental and theoretical approach. Indian J. Pure Appl. Phy. 2018, 56 (10), 814–829.

(21) Karpagakalyaani, G.; Magdaline, J. D.; Chithambarathanu, T.; Aruldhas, D.; Anuf, A. R., Spectroscopic (FT-IR, FT-Raman, NBO) investigation and molecular docking study of a herbicide compound Bifenox. Chem. Data Collect. 2020, 27, 100393.

(22) Meng, Y.; Yao, C.; Xue, S.; Yang, H., Application of Fourier transform infrared (FT-IR) spectroscopy in determination of microalgal compositions. Bioresour. Technol. 2014, 151, 347–354.

(23) Barroso-Bogeat, A.; Alexandre-Franco, M.; Fernández-González, C.; Gómez-Serrano, V., FT-IR Analysis of Pyrone and Chromene Structures in Activated Carbon. Energy. Fuels. 2014, 28 (6), 4096–4103.

(24) Lee, S.-G.; Na, D.; Park, C., Comparability of reference-based and reference-free transcriptome analysis approaches at the gene expression level. BMC Bioinformatics. 2021, 22, 1–9.

(25) Oiffer, T.; Leipold, F.; Süss, P.; Breite, D.; Griebel, J.; Khurram, M.; Branson, Y.; de Vries, E.; Schulze, A.; Helm, C. A.; Wei, R.; Bornscheuer, U. T., Chemo-Enzymatic Depolymerization of Functionalized Low-Molecular-Weight Polyethylene. Angew. Chem. Int. Ed. 2024, 63 (50), e202415012.

(26) Gambarini, V.; Pantos, O.; Kingsbury, J. M.; Weaver, L.; Handley, K. M.; Lear, G., PlasticDB: a database of microorganisms and proteins linked to plastic biodegradation. Database. 2022, 2022, baac008.

(27) Gan, Z.; Zhang, H., PMBD: a comprehensive plastics microbial biodegradation database. Database. 2019, 2019, baz119.

(28) Buchholz, P. C.; Feuerriegel, G.; Zhang, H.; Perez-Garcia, P.; Nover, L. L.; Chow, J.; Streit, W. R.; Pleiss, J., Plastics degradation by hydrolytic enzymes: The plastics-active enzymes database—PAZy. *Proteins: Struct., Funct.*, Bioinf. 2022, 90 (7), 1443–1456.

(29) Dong, J.; Fernández-Fueyo, E.; Hollmann, F.; Paul, C. E.; Pesic, M.; Schmidt, S.; Wang, Y.; Younes, S.; Zhang, W., Biocatalytic oxidation reactions: a chemist’s perspective. Angew. Chem. Int. Ed. 2018, 57 (30), 9238–9261.

(30) Gräff, M.; Buchholz, P. C. F.; Le Roes-Hill, M.; Pleiss, J., Multicopper oxidases: modular structure, sequence space, and evolutionary relationships. Proteins: Struct. Funct. Bioinf. 2020, 88 (10), 1329–1339.

(31) Teufel, F.; Almagro Armenteros, J. J.; Johansen, A. R.; Gíslason, M. H.; Pihl, S. I.; Tsirigos, K. D.; Winther, O.; Brunak, S.; von Heijne, G.; Nielsen, H., SignalP 6.0 predicts all five types of signal peptides using protein language models. Nat. Biotechnol. 2022, 40 (7), 1023–1025.

(32) Abramson, J.; Adler, J.; Dunger, J.; Evans, R.; Green, T.; Pritzel, A.; Ronneberger, O.; Willmore, L.; Ballard, A. J.; Bambrick, J.; Bodenstein, S. W.; Evans, D. A.; Hung, C.-C.; O’Neill, M.; Reiman, D.; Tunyasuvunakool, K.; Wu, Z.; Žemgulytė, A.; Arvaniti, E.; Beattie, C.; Bertolli, O.; Bridgland, A.; Cherepanov, A.; Congreve, M.; Cowen-Rivers, A. I.; Cowie, A.; Figurnov, M.; Fuchs, F. B.; Gladman, H.; Jain, R.; Khan, Y. A.; Low, C. M. R.; Perlin, K.; Potapenko, A.; Savy, P.; Singh, S.; Stecula, A.; Thillaisundaram, A.; Tong, C.; Yakneen, S.; Zhong, E. D.; Zielinski, M.; Žídek, A.; Bapst, V.; Kohli, P.; Jaderberg, M.; Hassabis, D.; Jumper, J. M., Accurate structure prediction of biomolecular interactions with AlphaFold 3. Nature. 2024, 630 (8016), 493–500.

(33) Quintanar, L.; Stoj, C.; Taylor, A. B.; Hart, P. J.; Kosman, D. J.; Solomon, E. I., Shall We Dance? How A Multicopper Oxidase Chooses Its Electron Transfer Partner. Acc. Chem. Res. 2007, 40 (6), 445–452.

(34) Melo, E. P.; Fernandes, A. T.; Durão, P.; Martins, L. O., Insight into stability of CotA laccase from the spore coat of *Bacillus subtilis*. Biochem. Soc. Trans. 2007, 35 (6), 1579–1582.

(35) Chovancova, E.; Pavelka, A.; Benes, P.; Strnad, O.; Brezovsky, J.; Kozlikova, B.; Gora, A.; Sustr, V.; Klvana, M.; Medek, P.; Biedermannova, L.; Sochor, J.; Damborsky, J., CAVER 3.0: A Tool for the Analysis of Transport Pathways in Dynamic Protein Structures. PLoS Comput. Biol. 2012, 8 (10), e1002708.

(36) Xie, T.; Liu, Z.; Wang, G., Structural basis for monolignol oxidation by a maize laccase. Nat. Plants. 2020, 6 (3), 231–237.

(37) Jin, J.; Arciszewski, J.; Auclair, K.; Jia, Z., Enzymatic polyethylene biorecycling: Confronting challenges and shaping the future. J. Hazard. Mater. 2023, 460, 132449.

(38) Martínez, A. T.; Speranza, M.; Ruiz-Dueñas, F. J.; Ferreira, P.; Camarero, S.; Guillén, F.; Martínez, M. J.; Gutiérrez, A.; del Río, J. C., Biodegradation of lignocellulosics: microbial, chemical, and enzymatic aspects of the fungal attack of lignin. Int. Microbiol. 2005, 8 (3), 195–204.

(39) Inderthal, H.; Tai, S. L.; Harrison, S. T. L., Non-Hydrolyzable Plastics – An Interdisciplinary Look at Plastic Bio-Oxidation. Trends Biotechnol. 2021, 39 (1), 12–23.

(40) Yao, Z.; Seong, H. J.; Jang, Y.-S., Environmental toxicity and decomposition of polyethylene. Ecotoxicol. Environ. Saf. 2022, 242, 113933.

(41) Gugumus, F., Re-examination of the thermal oxidation reactions of polymers 2. Thermal oxidation of polyethylene. Polym. Degrad. Stab. 2002, 76 (2), 329–340.

(42) Jiang, Q.; Li, Z.; Cui, Z.; Wei, R.; Nie, K.; Xu, H.; Liu, L., Quantum mechanical investigation of the oxidative cleavage of the C–C backbone bonds in polyethylene model molecules. Polymers. 2021, 13 (16), 2730.

(43) Jensen, R. K.; Korcek, S.; Mahoney, L. R.; Zinbo, M., Liquid-phase autoxidation of organic compounds at elevated temperatures. 1. The stirred flow reactor technique and analysis of primary products from n-hexadecane autoxidation at 120-180.degree.C. J. Am. Chem. Soc. 1979, 101 (25), 7574–7584.

(44) Jensen, R. K.; Korcek, S.; Mahoney, L. R.; Zinbo, M., Liquid-phase autoxidation of organic compounds at elevated temperatures. 2. Kinetics and mechanisms of the formation of cleavage products in n-hexadecane autoxidation. J. Am. Chem. Soc. 1981, 103 (7), 1742–1749.

(45) Jensen, R. K.; Korcek, S.; Zinbo, M., Formation, isomerization, and cyclization reactions of hydroperoxyalkyl radicals in hexadecane autoxidation at 160-190.degree.C. J. Am. Chem. Soc. 1992, 114 (20), 7742–7748.

(46) Wakayama, T.; Matsuhashi, H., Reaction of linear, branched, and cyclic alkanes catalyzed by Brönsted and Lewis acids on H-mordenite, H-beta, and sulfated zirconia. J. Mol. Catal. A Chem. 2005, 239 (1), 32–40.

(47) Guzik, M. W.; Nitkiewicz, T.; Wojnarowska, M.; Sołtysik, M.; Kenny, S. T.; Babu, R. P.; Best, M.; O’Connor, K. E., Robust process for high yield conversion of non-degradable polyethylene to a biodegradable plastic using a chemo-biotechnological approach. Waste Manag. 2021, 135, 60–69.

(48) Rabot, C.; Chen, Y.; Bijlani, S.; Chiang, Y. M.; Oakley, C. E.; Oakley, B. R.; Williams, T. J.; Wang, C. C., Conversion of polyethylenes into fungal secondary metabolites. Angew. Chem. 2023, 135 (4), e202214609.

(49) Sullivan, K. P.; Werner, A. Z.; Ramirez, K. J.; Ellis, L. D.; Bussard, J. R.; Black, B. A.; Brandner, D. G.; Bratti, F.; Buss, B. L.; Dong, X.; Haugen, S. J.; Ingraham, M. A.; Konev, M. O.; Michener, W. E.; Miscall, J.; Pardo, I.; Woodworth, S. P.; Guss, A. M.; Román-Leshkov, Y.; Stahl, S. S.; Beckham, G. T., Mixed plastics waste valorization through tandem chemical oxidation and biological funneling. Science. 2022, 378 (6616), 207–211.

(50) Vacher, M.; Fdez. Galván, I.; Ding, B.-W.; Schramm, S.; Berraud-Pache, R.; Naumov, P.; Ferré, N.; Liu, Y.-J.; Navizet, I.; Roca-Sanjuán, D.; Baader, W. J.; Lindh, R., Chemi- and Bioluminescence of Cyclic Peroxides. Chem. Rev. 2018, 118 (15), 6927–6974.

(51) Turro, N. J.; Lechtken, P.; Schore, N. E.; Schuster, G.; Steinmetzer, H. C.; Yekta, A., Tetramethyl-1,2-dioxetane. Experiments in chemiexcitation, chemiluminescence, photochemistry, chemical dynamics, and spectroscopy. Acc. Chem. Res. 1974, 7 (4), 97–105.

(52) Chen, Y.; Spiering, A. J. H.; Karthikeyan, S.; Peters, G. W. M.; Meijer, E. W.; Sijbesma, R. P., Mechanically induced chemiluminescence from polymers incorporating a 1,2-dioxetane unit in the main chain. Nat. Chem. 2012, 4 (7), 559–562.

(53) Bardají, D. K. R.; Furlan, J. P. R.; Stehling, E. G., Isolation of a polyethylene degrading *Paenibacillus* sp. from a landfill in Brazil. Arch. Microbiol. 2019, 201 (5), 699–704.

(54) Mathews, S. L.; Grunden, A. M.; Pawlak, J., Degradation of lignocellulose and lignin by *Paenibacillus glucanolyticus*. Int. Biodeterior. Biodegrad. 2016, 110, 79–86.

(55) Govarthanan, M.; Mythili, R.; Selvankumar, T.; Kamala-Kannan, S.; Rajasekar, A.; Chang, Y.-C., Bioremediation of heavy metals using an endophytic bacterium *Paenibacillus* sp. RM isolated from the roots of Tridax procumbens. 3 Biotech. 2016, 6 (2), 242.

(56) Nassar, H. N.; Abu Amr, S. S.; El-Gendy, N. S., Biodesulfurization of refractory sulfur compounds in petro-diesel by a novel hydrocarbon tolerable strain *Paenibacillus glucanolyticus* HN4. Environ. Sci. Pollut. Res. 2021, 28 (7), 8102–8116.

(57) Nagarathna, S. V.; Chandramouli Swamy, T. M.; Reddy, P. V.; Kanade, S. R.; Nayak, A. S., Metabolism of Benzo[a]pyrene by *Paenibacillus* sp. PRNK-6 through novel metabolite phenalene-1,9-dicarboxylic acid. Int. Biodeterior. Biodegrad. 2025, 196, 105938.

(58) Park, S. Y.; Kim, C. G., Biodegradation of micro-polyethylene particles by bacterial colonization of a mixed microbial consortium isolated from a landfill site. Chemosphere. 2019, 222, 527–533.

(59) Chin, C.-S.; Alexander, D. H.; Marks, P.; Klammer, A. A.; Drake, J.; Heiner, C.; Clum, A.; Copeland, A.; Huddleston, J.; Eichler, E. E.; Turner, S. W.; Korlach, J., Nonhybrid, finished microbial genome assemblies from long-read SMRT sequencing data. Nat. Methods. 2013, 10 (6), 563–569.

(60) Bolger, A. M.; Lohse, M.; Usadel, B., Trimmomatic: a flexible trimmer for Illumina sequence data. Bioinformatics. 2014, 30 (15), 2114–2120.

(61) Walker, B. J.; Abeel, T.; Shea, T.; Priest, M.; Abouelliel, A.; Sakthikumar, S.; Cuomo, C. A.; Zeng, Q.; Wortman, J.; Young, S. K., Pilon: an integrated tool for comprehensive microbial variant detection and genome assembly improvement. PloS One. 2014, 9 (11), e112963.

(62) Seemann, T., Prokka: rapid prokaryotic genome annotation. Bioinformatics 2014, 30 (14), 2068–2069.

(63) Grant, J. R.; Enns, E.; Marinier, E.; Mandal, A.; Herman, E. K.; Chen, C.-y.; Graham, M.; Van Domselaar, G.; Stothard, P., Proksee: in-depth characterization and visualization of bacterial genomes. Nucleic Acids Research. 2023, 51 (W1), W484–W492.

(64) Langmead, B.; Wilks, C.; Antonescu, V.; Charles, R., Scaling read aligners to hundreds of threads on general-purpose processors. Bioinformatics 2018, 35 (3), 421–432.

(65) Shumate, A.; Wong, B.; Pertea, G.; Pertea, M., Improved transcriptome assembly using a hybrid of long and short reads with StringTie. PLoS Comput. Biol. 2022, 18 (6), e1009730.

(66) Robinson, M. D.; McCarthy, D. J.; Smyth, G. K., edgeR: a Bioconductor package for differential expression analysis of digital gene expression data. Bioinformatics. 2010, 26 (1), 139–140.

(67) Li, L.; Stoeckert, C. J.; Roos, D. S., OrthoMCL: identification of ortholog groups for eukaryotic genomes. Genome Res. 2003, 13 (9), 2178–2189.

(68) Katoh, K.; Misawa, K.; Kuma, K. i.; Miyata, T., MAFFT: a novel method for rapid multiple sequence alignment based on fast Fourier transform. Nucleic Acids Research. 2002, 30 (14), 3059–3066.

(69) Capella-Gutiérrez, S.; Silla-Martínez, J. M.; Gabaldón, T., trimAl: a tool for automated alignment trimming in large-scale phylogenetic analyses. Bioinformatics. 2009, 25 (15), 1972–1973.

(70) Kalyaanamoorthy, S.; Minh, B. Q.; Wong, T. K. F.; von Haeseler, A.; Jermiin, L. S., ModelFinder: fast model selection for accurate phylogenetic estimates. Nat. Methods. 2017, 14 (6), 587–589.

(71) Minh, B. Q.; Schmidt, H. A.; Chernomor, O.; Schrempf, D.; Woodhams, M. D.; von Haeseler, A.; Lanfear, R., IQ-TREE 2: New Models and Efficient Methods for Phylogenetic Inference in the Genomic Era. Mol. Biol. Evol. 2020, 37 (5), 1530–1534.

(72) Egbewale, S. O.; Kumar, A.; Mokoena, M. P.; Olaniran, A. O., Purification, characterization and three-dimensional structure prediction of multicopper oxidase Laccases from *Trichoderma lixii* FLU1 and *Talaromyces pinophilus* FLU12. Sci. Rep. 2024, 14 (1), 13371.

(73) Bourbonnais, R.; Paice, M. G., Oxidation of non-phenolic substrates: An expanded role for laccase in lignin biodegradation. FEBS Lett. 1990, 267 (1), 99–102.

(74) Leonowicz, A.; Grzywnowicz, K., Quantitative estimation of laccase forms in some white-rot fungi using syringaldazine as a substrate. Enzyme Microb. Technol. 1981, 3 (1), 55–58.

