## supporting information for "A Multicopper Oxidase from *Paenibacillus polyethylenelyticus* JNU01 Oxidizes Polyethylene"

**This Supporting information file includes:**

Figures S1 to S9

Tables S1 to S5

**Other Supplementary data for this manuscript include the following:**

Supplementary data S1 to S5

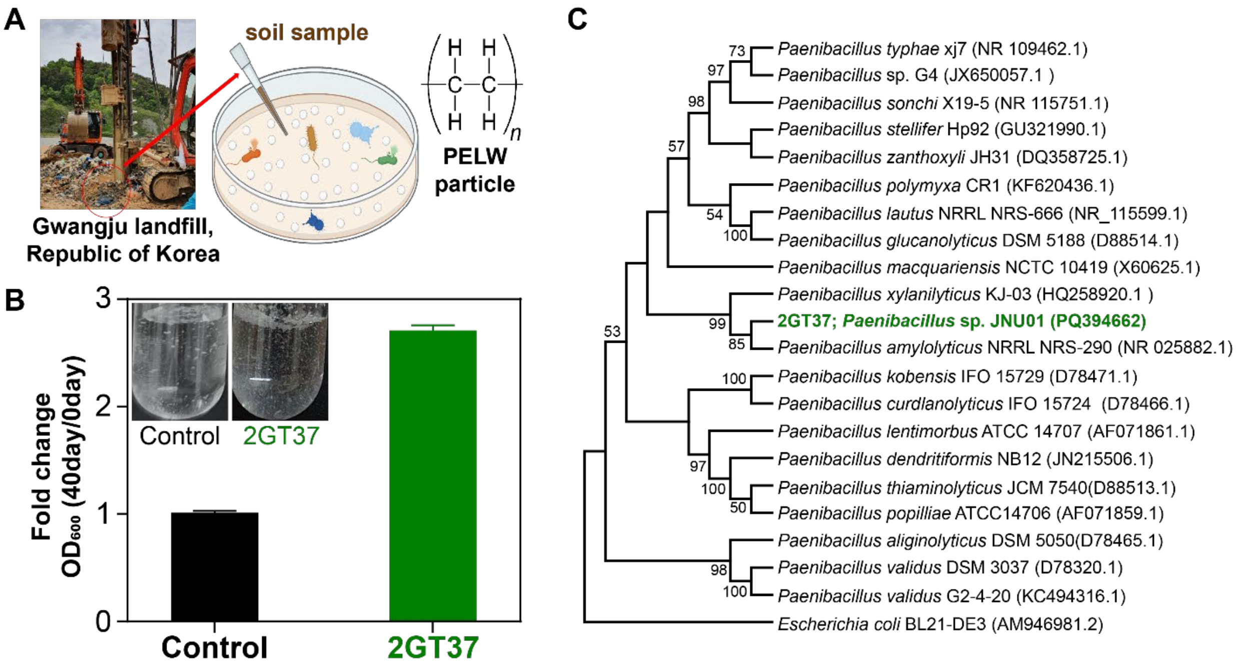


**Figure S1. Identification process for PELW biodegrading strains.** (**A**) The screening of PELW-biodegrading strain derived from Gwangju Landfill. (**B**) The fold changes of day 40 ($OD_{600}$)/0 DAY($OD_{600}$) for strain 2GT37 in liquid M9 minimal broth supplemented with 1 g/L of PELW powder ($n = 2$, independent experiments). (**C**) A phylogenetic tree based on strain 2GT37 (*Paenibacillus* sp. JNU01) and other bacteria's 16s rRNA sequences.

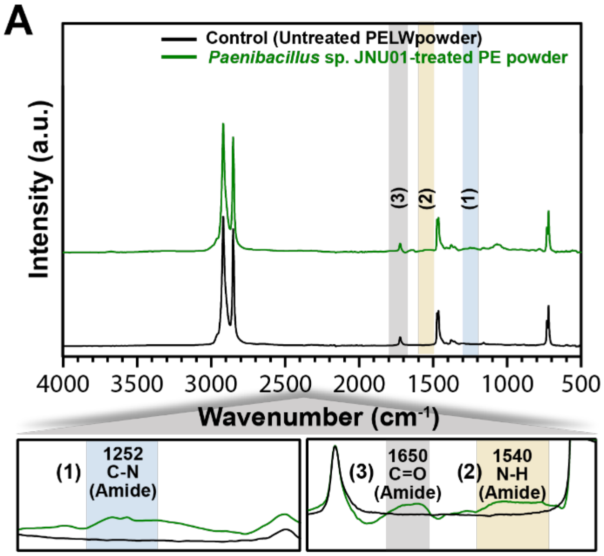


**Figure S2. Effect of extended washing on FT-IR spectra of *Paenibacillus polyethylenelyticus* JNU01–treated PELW powders.** (**A**) Spectrum of the powder washed only with methanol (4 times), in which amide-related peaks (1252$^{-1}$ (peak 1), 1540$^{-1}$ (peak 2), and 1650 cm$^{-1}$ (peak 3)) are observed. (**B**) Spectrum of the powder further washed with 1M NaOH (5 times) and distilled water (5 times), in which the amide-related peaks are removed.

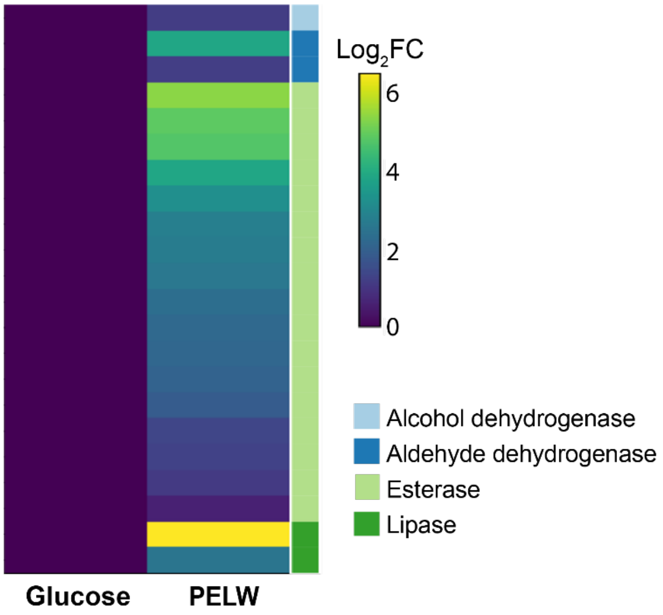


**Figure S3. A heatmap shows 22 potential candidate genes for PELW biodegradation.** The $Log_2FC$ ($log_2$-transformed fold change) data were calculated using edgeR. Each gene is color coded based on its enzyme class.

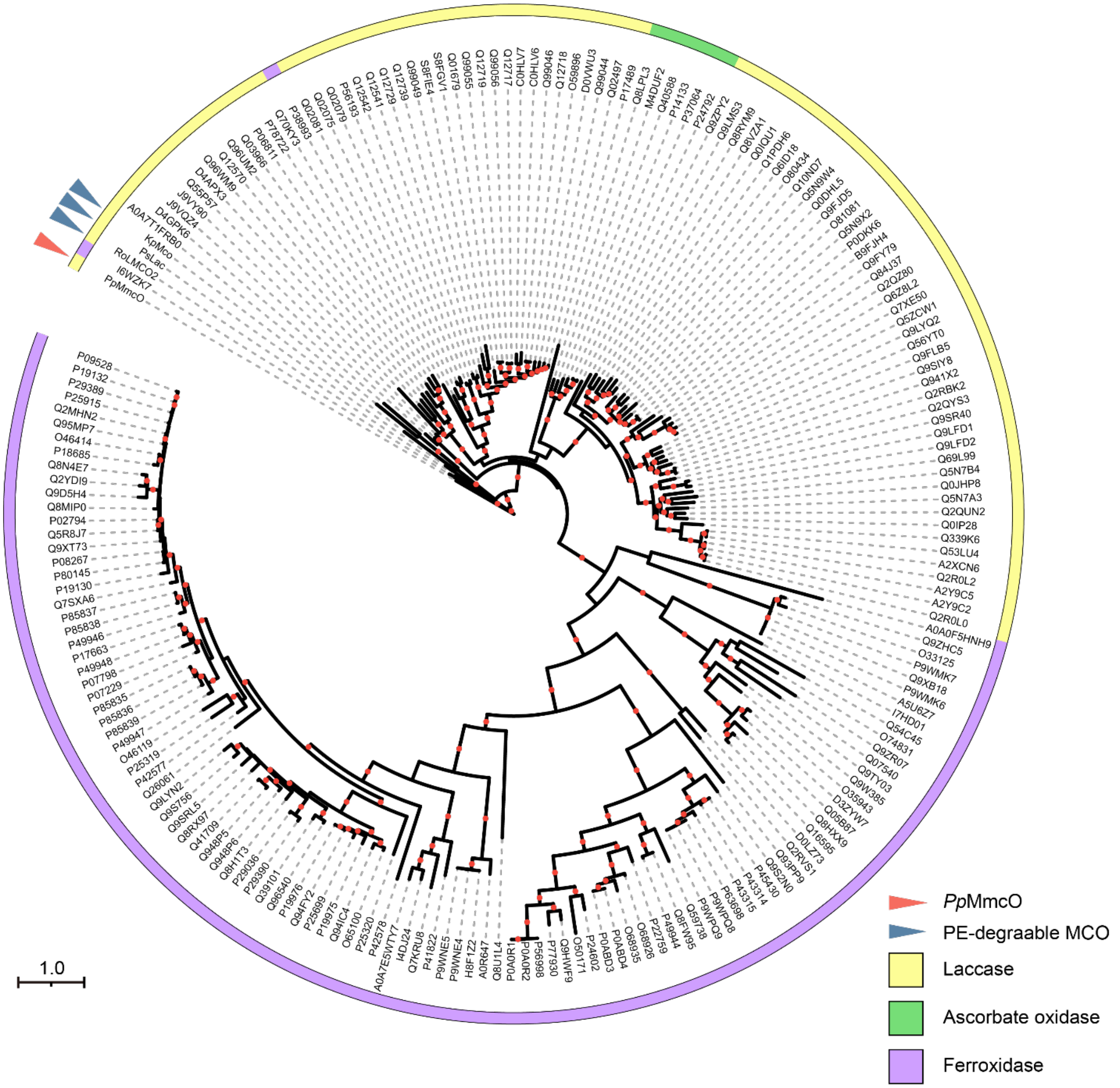


**Figure S4. Phylogenetic tree of *Pp*MmcO$^{WT}$. A phylogenetic tree of *Pp*MmcO with other multicopper oxidases (MCOs).** The phylogenetic tree shows *Pp*MmcO along with 83 laccases, 6 ascorbate oxidases, 109 ferroxidases, and 3 PE-degradable MCOs. The organism sources of the proteins and their accession numbers are shown at the leaf nodes. The MCO families are distinguished by different colors. Red arrows indicate *Pp*MmcO, and blue arrows indicate PE-degradable MCOs. Bootstrap values greater than 80 are shown at the midpoints of the branches. Scale bar: 1.0 amino acid substitutions per site.

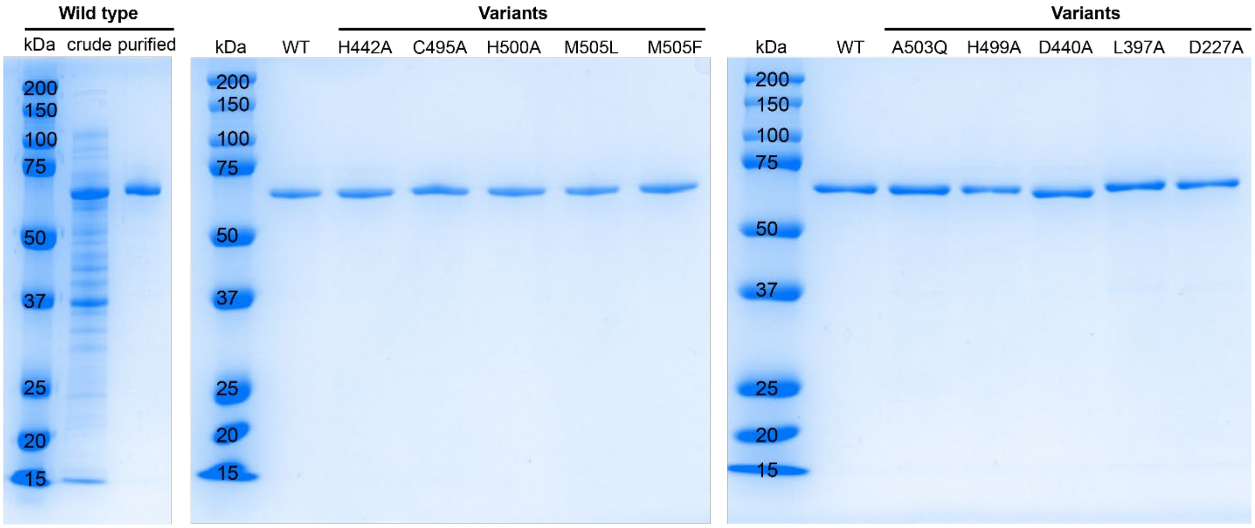


**Figure S5. SDS-PAGE results for the *Pp*MmcO$^{WT}$ and its variants.** The molecular weights of the *Pp*MmcO$^{WT}$ and its variants are approximately 60 kDa, as determined by SDS-PAGE.

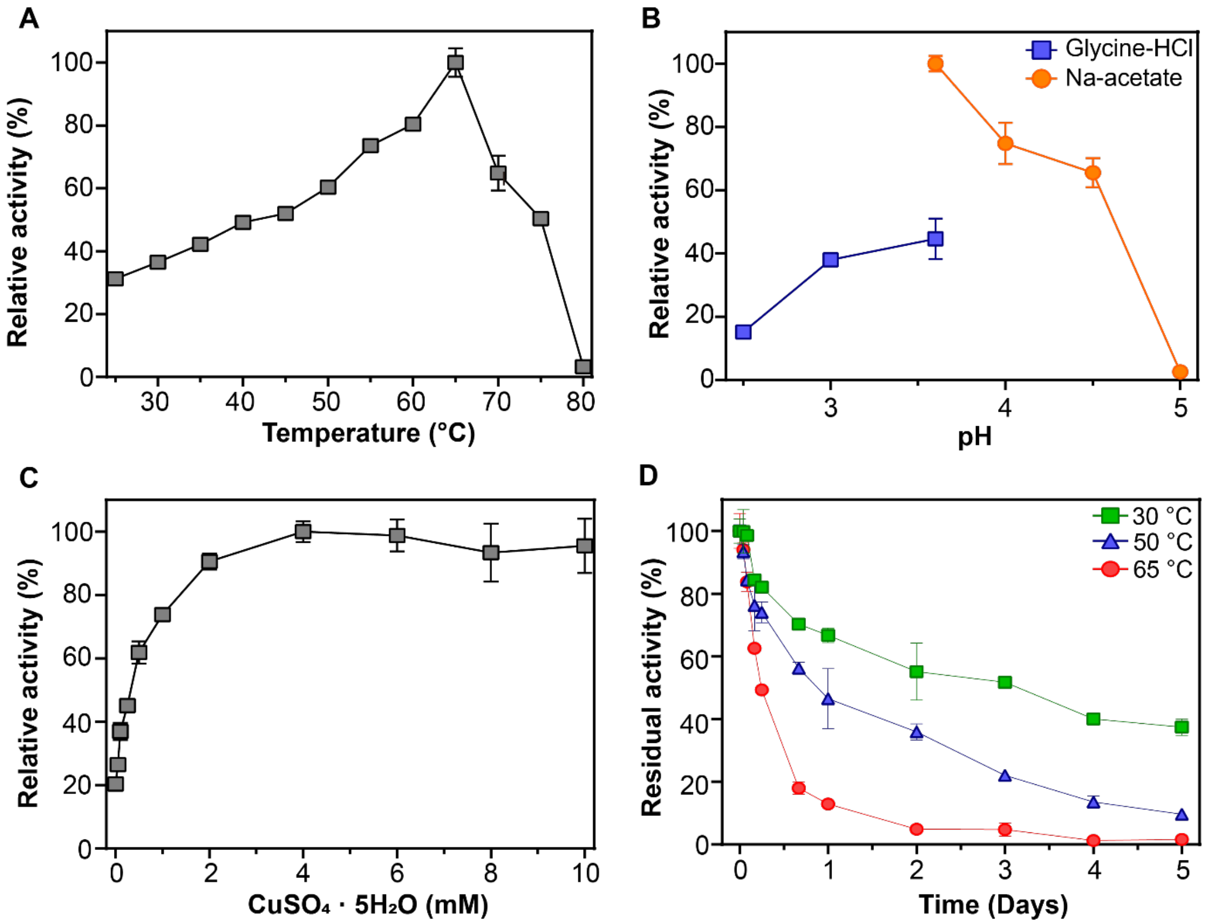


**Figure S6. Optimization of *Pp*MmcO$^{WT}$.** (**A**) Temperature: reactions were conducted for 2 min in 20 mM Na-acetate buffer (pH 4.0) with 2.0 mM $CuSO_4$, 0.50 mM ABTS, and 0.010 mg/mL of enzyme. (**B**) pH: reactions were conducted for 2 min at 65 °C in 20 mM Glycine-HCl or Na-acetate buffer with 2.0 mM $CuSO_4$, 0.50 mM ABTS, and 0.010 mg/mL of enzyme. (**C**) $CuSO_4$ concentration: reactions were conducted for 2 min at 65 °C in 20 mM Na-acetate buffer (pH 3.6) with 0–10 mM $CuSO_4$, 0.50 mM ABTS, and 0.010 mg/mL enzyme. (**D**) Thermal stability: residual activity was measured over 5 d at 30 °C, 50 °C, and 65 °C in 20 mM Na-acetate buffer (pH 3.6) with 2.0 mM $CuSO_4$, 0.50 mM ABTS, and 0.010 mg/mL enzyme. All experiments were performed in duplicate ($n = 2$, independent experiments).

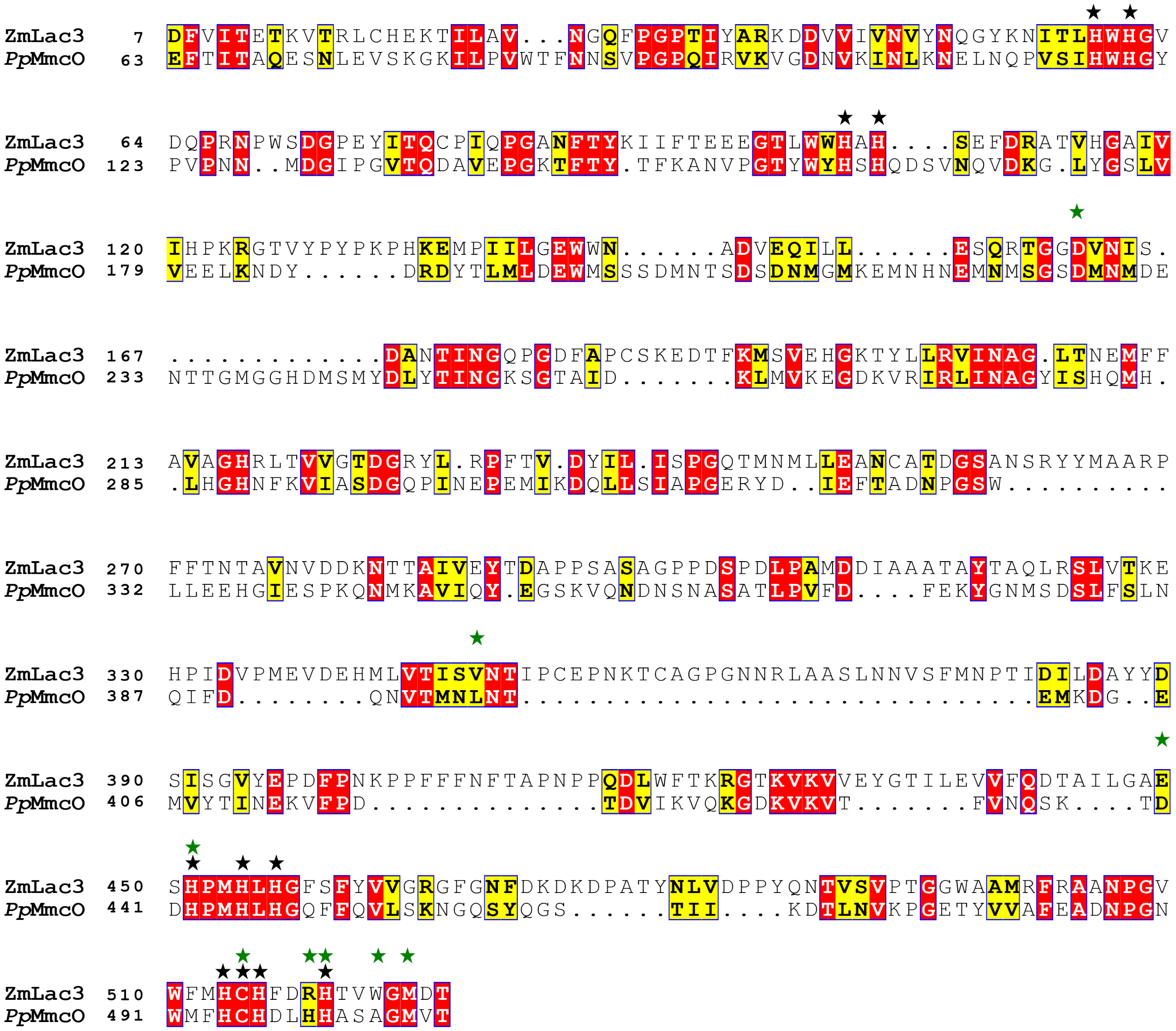


**Figure S7. Amino acid alignment of ZmLac3 and *Pp*MmcO$^{WT}$.** Black stars indicate highly conserved residues around copper atoms and green stars denote residues selected as variants around the T1 copper atom.

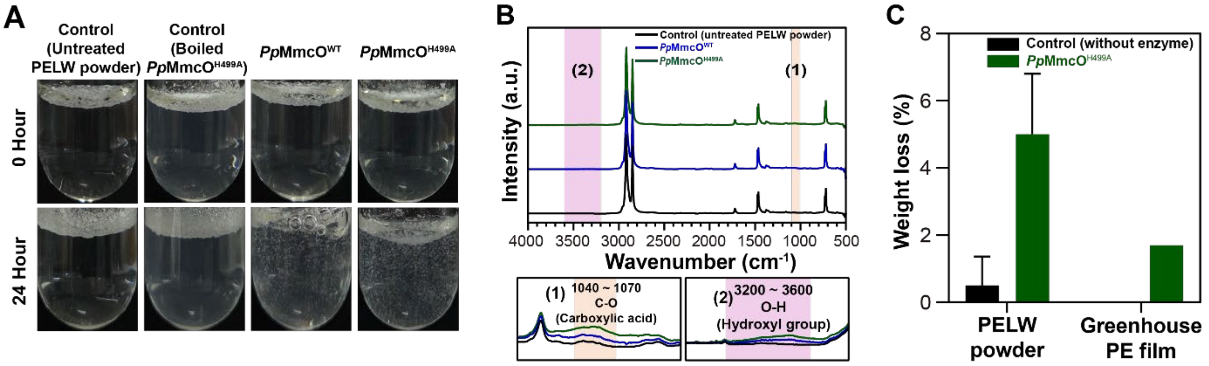


**Figure S8. Structural changes in PELW powder and weight-loss analysis of PELW and greenhouse PE films after *Pp*MmcO$^{H499A}$ treatment.** (**A**) Images of the control solutions (untreated PELW powder without enzyme and boiled *Pp*MmcO$^{H499A}$and the enzyme reaction solutions containing *Pp*MmcO$^{WT}$ and *Pp*MmcO$^{H499A}$ at 0 and 24 h. (**B**) FT-IR spectra of control (untreated PELW powder), *Pp*MmcO$^{WT}$-treated, and *Pp*MmcO$^{H499A}$-treated PELW powders. (**C**) Weight loss (%) of the solution-cast PELW film and greenhouse PE film after treatment with *Pp*MmcO$^{H499A}$. The control sample was incubated under identical conditions without *Pp*MmcO$^{H499A}$ (n = 3, mean ± s.d.)

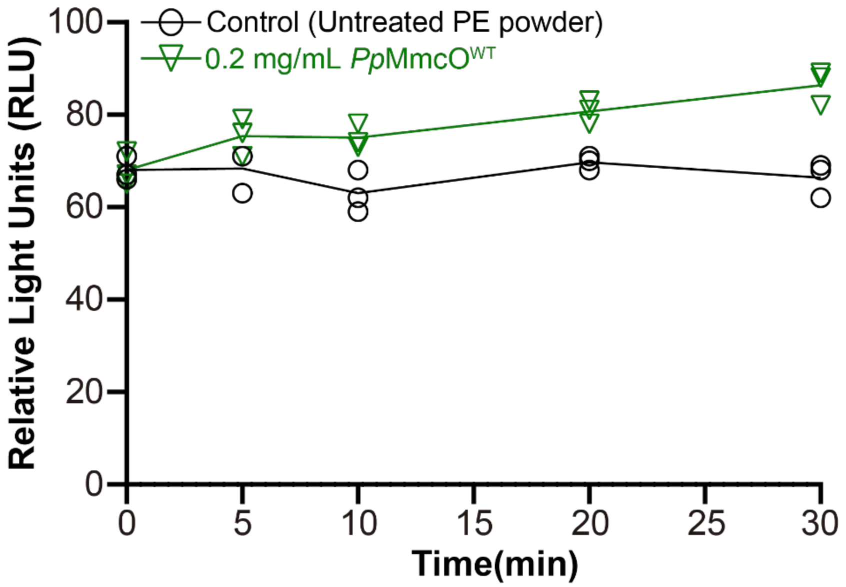


**Figure S9. Luminescence measurement of *Pp*MmcO$^{WT}$.** The luminescence intensity ($n = 3$ independent experiments; mean ± s.d.) of the reaction solution containing 0.2 mg/mL *Pp*MmcO$^{WT}$ and PELW powder was measured over time, along with the control solution containing untreated PELW powder.

**Table S1. Identification of metabolites from GC-MS analysis of *Paenibacillus* sp. JNU01 cultured in PELW media.**

| Number | min | Similarity (%) | Compound name | Molecular weight | Formula |
|---|---|---|---|---|---|
| 1 | 5.32 | 94 | 3,4,5-Trimethylheptane | 142 | $C_{10}H_{22}$ |
| 2 | 5.5 | 93 | 2,3,3-Trimethylheptane | 128 | $C_9H_{20}$ |
| 3 | 12.01 | 93 | 3-Ethyl-3-methylheptane | 142 | $C_{10}H_{22}$ |
| 4 | 13.26 | 92 | *n*-Dodecane | 170 | $C_{12}H_{26}$ |
| 5 | 17.87 | 91 | 5-Methyl-5-propylnonane | 185 | $C_{13}H_{28}$ |
| 6 | 19.47 | 91 | *n*-Tetradecane | 198 | $C_{14}H_{30}$ |
| 7 | 23.88 | 91 | *n*-Hexadecane | 226 | $C_{16}H_{34}$ |
| 8 | 26.08 | 83 | 11-Oxododecanoic acid | 214 | $C_{12}H_{22}O_3$ |
| 9 | 28.15 | 84 | *n*-Eicosane | 282 | $C_{20}H_{42}$ |
| 10 | 28.4 | 86 | Eicosanoic acid | 312 | $C_{20}H_{40}O_2$ |
| 11 | 35.92 | 82 | 1-Tricosanol | 340 | $C_{23}H_{48}O$ |

### Table S2. Primer sequences used in study.

| Cloning | | Primer sequence |
|---|---|---|
| *Pp*MmcO Wild-Type | Insert | Forward: 5′-TGTATTTTCAGGGCCATATGTGTACTAATGCGAT-3′ |
| | | Reverse: 5′-CCTTTGAATTCCGGATCTTATTCCGCTACGTTCT-3′ |
| | vector | Forward: 5′-AGAACGTAGCGGAATAAGATCCGGAATTCAAAGG-3′ |
| | | Reverse: 5′-ATCGCATTAGTACACATATGGCCCTGAAAATACA-3′ |
| **Site-directed mutagenesis** | | **Primer sequence** |
| *Pp*MmcO Variants | H442A | Forward: 5′-CAGTCCAAAACGGATGACGCTCCTATGCATCTCCATGG-3′ |
| | | Reverse: 5′-CCATGGAGATGCATAGGAGCGTCATCCGTTTTGGACTG-3′ |
| | C495A | Forward: 5′-CCTGGAAATTGGATGTTCCATGCCCATGACTTGCATCATGC-3′ |
| | | Reverse: 5′-GCATGATGCAAGTCATGGGCATGGAACATCCAATTTCCAGG-3′ |
| | H500A | Forward: 5′-TTCCATTGCCATGACTTGCATGCTGCTTCCGCGGG-3′ |
| | | Reverse: 5′-CCCGCGGAAGCAGCATGCAAGTCATGGCAATGGAA-3′ |
| | M505L | Forward: 5′-CATCATGCTTCCGCGGGATTGGTAACGGAAGT-3′ |
| | | Reverse: 5′-ACTTCCGTTACCAATCCCGCGGAAGCATGATG-3′ |
| | M505F | Forward: 5′-CATCATGCTTCCGCGGGATTCGTAACGGAAGTCATGTAT-3′ |
| | | Reverse: 5′-ATACATGACTTCCGTTACGAATCCCGCGGAAGCATGATG-3′ |
| | A503Q | Forward: 5′-CTTGCATCATGCTTCCCAGGGAATGGTAACGGAAG-3′ |
| | | Reverse: 5′-CTTCCGTTACCATTCCCTGGGAAGCATGATGCAAG-3′ |
| | H499A | Forward: 5′-CATTGCCATGACTTGGCTCATGCTTCCGCGGG-3′ |
| | | Reverse: 5′-CCCGCGGAAGCATGAGCCAAGTCATGGCAATG-3′ |
| | D440A | Forward: 5′-GAATCAGTCCAAAACGGCTGACCATCCTATGCATC-3′ |
| | | Reverse: 5′-GATGCATAGGATGGTCAGCCGTTTTGGACTGATTC-3′ |
| | L397A | Forward: 5′-AACGTGACAATGAATGCAAATACAGAAATGAAAG-3′ |
| | | Reverse: 5′-CTTTCATTTCTGTATTTGCATTCATTGTCACGTT-3′ |
| | D227A | Forward: 5′-GAATATGTCCGGATCGGCTATGAACATGGACGAG-3′ |
| | | Reverse: 5′-CTCGTCCATGTTCATAGCCGATCCGGACATATTC-3′ |

**Table S3. Amino acid sequences of *P. polyethylenelyticus* JNU01's wild-type (WT) *Pp*MmcO and its variants.** Signal peptide amino acid sequences are shown in red, and the mutated amino acid residues are indicated in blue.

| Target Enzyme | Amino acid sequence |
|---|---|
| *Pp*MmcO$^{WT}$ | MKLMLLKRFIVIGSLAILATGCTNAIKNNAMEGMDHSNTNNPT SAGSSPVKFTSSTPVVNGKEFTITAQESNLEVSKGKILPVWTFNN SVPGPQIRVKVGDNVKINLKNELNQPVSIHWHGYPVPNNMDGI PGVTQDAVEPGKTFTYTFKANVPGTYWYHSHQDSVNQVDKGL YGSLVVEELKNDYDRDYTLMLDEWMSSSDMNTSDSDNMGMK EMNHNEMNMSGSDMNMDENTTGMGGHDMSMYDLYTINGKS GTAIDKLMVKEGDKVRIRLINAGYISHQMHLHGHNFKVIASDG QPINEPEMIKDQLLSIAPGERYDIEFTADNPGSWLLEEHGIESPKQ NMKAVIQYEGSKVQNDNSNASATLPVFDFEKYGNMSDSLFSLN QIFDQNVTMNLNTEMKDGEMVYTINEKVFPDTDVIKVQKGDK VKVTFVNQSKTDDHPMHLHGQFFQVLSKNGQSYQGSTIIKDTL NVKPGETYVVAFEADNPGNWMFHCHDLHHASAGMVTEVMY TDYESNYTPDPSVENVAE |

**Table S4. List of compounds detected through GC-MS analysis of enzyme reaction solution of *Pp*MmcO wild-type and PE powder.** Compound identified in the chromatogram in Fig. 4.

| Number | min | Compound Similarity (%) by Enzyme Concentration | | | Compound name | Molecular weight | Formula |
|---|---|---|---|---|---|---|---|
| | | 0.2 mg/mL | 1.0 mg/mL | 3.0 mg/mL | | | |
| 1 | 32.79 | 87 | 85 | 86 | 2-Heptadecanone | 254 | $C_{17}H_{34}O$ |
| 2 | 34.41 | 92 | 92 | 92 | *n*-Eicosane | 282 | $C_{20}H_{42}$ |
| 3 | 34.82 | 82 | 85 | 87 | *cis*-10-Heptadecenoic acid trimethylsilyl ester | 340 | $C_{20}H_{40}O_2Si$ |
| 4 | 36.08 | 92 | 92 | 92 | 2-Nonadecanone | 282 | $C_{19}H_{38}O$ |
| 5 | 36.3 | 88 | 90 | 92 | *trans*-9-Octadecenoic acid trimethylsilyl ester | 354 | $C_{21}H_{42}O_2Si$ |
| 6 | 37.52 | 91 | 91 | 91 | 2-Methylpentacosane | 366 | $C_{26}H_{54}$ |
| 7 | 37.9 | 83 | 83 | 85 | *cis*-10-Nonadecenoic acid trimethylsilyl ester | 368 | $C_{22}H_{44}O_2Si$ |
| 8 | 39.11 | 86 | 86 | 86 | 2-Nonacosanone | 422 | $C_{29}H_{58}O$ |
| 9 | 40.38 | 92 | 92 | 92 | *n*-Triacontane | 422 | $C_{30}H_{62}$ |
| 10 | 41.9 | 91 | 88 | 89 | 2-Tritriacontanone | 478 | $C_{33}H_{66}O$ |

**Table S5. List of compounds detected through GC-MS analysis of enzyme reaction solutions of *Pp*MmcO wild-type and the variant (H449A) with PELW powder.** Compound identified in the chromatogram in Fig. 6E.

| Number | min | Compound Similarity (%) for Wild-Type and the Variant (H449A) | | Compound name | Molecular weight | Formula |
|---|---|---|---|---|---|---|
| | | Wild-type | H449A | | | |
| 1 | 32.9 | 83 | 84 | 2-Heptadecanone | 254 | $C_{17}H_{34}O$ |
| 2 | 34.52 | 92 | 93 | *n*-Eicosane | 282 | $C_{20}H_{42}$ |
| 3 | 34.92 | 81 | 82 | *cis*-10-Heptadecenoic acid trimethylsilyl ester | 340 | $C_{20}H_{40}O_2Si$ |
| 4 | 36.2 | 84 | 86 | 2-Nonadecanone | 282 | $C_{19}H_{38}O$ |
| 5 | 36.41 | 85 | 88 | *trans*-9-Octadecenoic acid trimethylsilyl ester | 354 | $C_{21}H_{42}O_2Si$ |
| 6 | 37.63 | 90 | 91 | 2-Methylpentacosane | 366 | $C_{26}H_{54}$ |
| 7 | 38 | 81 | 81 | *cis*-10-Nonadecenoic acid trimethylsilyl ester | 368 | $C_{22}H_{44}O_2Si$ |
| 8 | 39.22 | 88 | 89 | 2-Nonacosanone | 422 | $C_{29}H_{58}O$ |
| 9 | 40.5 | 87 | 87 | *n*-Triacontane | 422 | $C_{30}H_{62}$ |
| 10 | 42 | 89 | 91 | 2-Tritriacontanone | 478 | $C_{33}H_{66}O$ |

**Supplementary data S1. (separate file)**

Average nucleotide identity (ANI) among species and strains.

**Supplementary data S2. (separate file)**

RNA-Seq analysis statistics.

**Supplementary data S3. (separate file)**

A list of 343 upregulated DEGs in PELW.

**Supplementary data S4. (separate file)**

A list of 798 downregulated DEGs in PELW.

**Supplementary data S5. (separate file)**

The amino acid sequences of enzymes used in the phylogenetic tree of *Pp*MmcO (Fig. S4).
